# Increased susceptibility to intracellular bacteria and necrotic inflammation driven by a dysregulated macrophage response to TNF

**DOI:** 10.1101/238873

**Authors:** Bidisha Bhattacharya, Sujoy Chattrerjee, Robert Berland, Alexander Pichugin, Yuanwei Gao, John Connor, Alexander Ivanov, Bo-Shiun Yan, Lester Kobzik, Igor Kramnik

**Affiliations:** The National Emerging Infectious Diseases Laboratory; Department of Medicine, Pulmonary Center; Department of Microbiology Boston University School of Medicine 620 Albany Street, Boston, MA 02118; Department of Chemistry & Chemical Biology, Northeastern University; Institute of Biochemistry and Molecular Biology, National Taiwan University Medical College, Taipei, Taiwan; Harvard T. H. Chan School of Public Health; Department of Integrative Physiology and Pathobiology, Tufts University School of Medicine; Department of Cellular Immunology, Malaria Vaccine Branch, Walter Reed Army Institute of Research, 503 Robert Grant Ave, Silver Spring, MD 20910; Department of Pharmacokinetics, Pharmacodynamics & Drug Metabolism (PPDM), Merck 770 Sumneytown Pike, West Point, PA 19446

## Abstract

Host susceptibility to tuberculosis and several other intracellular bacteria is controlled by a mouse genetic locus, sst1. Necrotic inflammatory lesions, similar to human TB granulomas, are a hallmark of the sst1 susceptible phenotype. Our previous work established that increased disease severity in sst1 susceptible mice reflects dysfunctional macrophage effector or tolerance mechanisms, but the molecular mechanisms have been unclear.

We demonstrate that *sst1-*deficient macrophages develop aberrant, biphasic responses to TNF, characterized by super-induction of stress and type I interferon pathways after prolonged TNF stimulation. This late stage response was initiated by oxidative stress and Myc. It was driven via a JNK - IFNβ - PKR feed-forward circuit locking the susceptible macrophages in a state of escalating stress. Consequently, prolonged TNF stimulation of the susceptible macrophages reduced their resilience to subsequent infection with intracellular bacteria.

The data support a generalizable paradigm in host - pathogen interactions, where susceptibility emerges gradually within inflammatory tissue due to imbalanced macrophage responses to growth, differentiation, activation and stress stimuli prior to encountering pathogens. This explains how successful pathogens may locally bypass mechanisms of resistance in otherwise immunocompetent hosts and suggests novel therapeutic strategies.

## INTRODUCTION

Susceptibility to infectious agents varies considerably within populations of natural hosts, manifest as a spectrum of permissiveness to individual pathogens, and a range of severity and outcomes of infectious diseases. These heterogeneous outcomes are governed by multi-dimensional interactions of genetic, behavioral and environmental factors. A corollary is that key pathogenic properties of microbes are best revealed within the specific context of susceptible hosts. Ultimately, understanding general and specific mechanisms of host susceptibility to individual pathogens or groups of pathogens is vital for the development of effective preventive and therapeutic strategies, including against infectious diseases caused by antibiotic-resistant bacteria.

Susceptibility can be either constitutive, due to genetic variation[1], or acquired, due to co-infections, malnutrition, stress, ageing and other factors, causing a functional decline in host immune and homeostatic systems. Less understood, are intrinsic mechanisms of susceptibility that develop within specific tissue environments. This type of induced susceptibility may develop in organ-specific manner in otherwise immune-competent individuals. For example, *Mycobacterium tuberculosis* (Mtb) targets lungs of immune-competent hosts to ensure its survival in granulomas and later transmission via aerosols[2]. The acquisition of permissiveness to infection by certain host cells in TB granulomas does not equate to loss of control mechanisms at the whole organism level[3, 4]. Thus, induced susceptibility may be effectively exploited by specialized pathogens, and, play an important role in their transmission and coevolution with hosts.

So far, susceptibility to infection has been largely viewed as an innate or induced defect of host immunity[5]. However, host susceptibility is not simply a failure of an essential pathogen recognition or effector mechanisms. In broad terms susceptibility to infectious diseases is determined by interactions of at least two groups of factors: 1) an array of effector mechanisms that counter pathogen fitness, i.e. ability to survive and replicate within the host tissues, and 2) host tolerance to infection defined at the whole organism level as host ability to maintain a certain health status in the presence of substantial pathogenic loads[6-8]. Successful pathogens have been shown to avoid or suppress mechanisms of host resistance. How they interact with host tolerance is less well understood, because current knowledge of physiological and genetic basis of host tolerance is limited [9, 10].

At the tissue level tolerance to infection is determined by cell resilience to stressors within inflammatory milieu induced by pathogens’ invasion. Indeed, mechanisms essential for host resistance to many bacterial infections, such as reactive oxygen and nitrogen species, may also induce collateral tissue damage, unless they are countered by host mechanisms of stress resilience. The salient feature of host tolerance strategies, as opposed to resistance, is their focus on maintaining host cell and tissue homeostasis [11, 12], and, therefore, tolerance mechanisms are expected to operate in pathogenesis-specific rather than pathogen-specific manner. Therefore, their failure may increase tissue damage and severity of infectious diseases caused by taxonomically unrelated agents.

To study mechanisms of host susceptibility to virulent Mtb, we developed a mouse model of human-like necrotic TB granulomas using C3HeB/FeJ (C3H) mice[13-15]. Using forward genetic analysis in a cross of the C3H with relatively TB resistant C57BL/6J (B6) mice, we have identified a novel genetic locus *sst1* (***<u>s</u>***uper***<u>s</u>***usceptibility to ***t***uberculosis ***<u>1</u>***), as a specific determinant of necrosis in TB granulomas[16]. Congenic mice that carry the C3HeB/FeJ-derived *sst1* susceptibility allele (sst1S) on the resistant B6 background, B6-*sst1^C3H,S^*(B6-sst1S), developed large, well-organized pulmonary necrotic granulomas after a low dose aerosol infection, even though initially these mice controlled Mtb replication similarly to the parental B6 mice[17]. The necrosis in the B6-sst1S TB lesions occurred with bacterial loads approximately 50-fold lower than in the parental C3HeB/FeJ mice indicating that extreme bacterial loads did not drive the sst1S-mediated necrosis. These data demonstrated for the first time that mechanisms controlling Mtb-inflicted necrotic damage could be genetically uncoupled from the effector immune mechanisms controlling the bacterial load.

Subsequent experiments using the *sst1* congenic mice revealed that the *sst1* locus controlled macrophage interactions with Mtb and other intracellular bacterial pathogens – *Listeria monocytogenes*[18] and *Chlamydia pneumonia*[19]. Using positional cloning approach, we have identified a variant of the interferon-inducible nuclear protein Sp110, Ipr1 (intracellular pathogen resistance 1), as a strong candidate gene, whose expression is completely abolished in sst1S macrophages[20]. The human homologue of Ipr1 - Sp110b - has been shown to play important roles in immunity to infections[21] and in control of macrophage activation[22]. However, the molecular mechanisms explaining the role of *sst1/Sp110* in the pathogenesis of TB and other infections remain to be elucidated.

Here we present evidence that the mouse genetic locus *sst1* is a broad determinant of host tolerance that controls macrophage resilience to stress induced by TNF, a major cytokine required for innate immune responses to many infectious agents, including Mtb [23]. Compared to the wild type, TNF stimulation of the sst1S congenic macrophages induced a complex cascade beginning with proteotoxic stress, Myc hyperactivity and super-induction of type I interferon (IFN-I), and culminating with an escalating integrated stress response (ISR). We delineated the hierarchy and mechanisms driving this cascade and demonstrated its role in induced susceptibility of macrophages to yet another intracellular bacterial pathogen, *Francisella tularensis* LVS, in vitro and in vivo. These findings delineate a common mechanism of inflammation-induced susceptibility to several taxonomically unrelated intracellular bacterial pathogens induced by a dysregulated macrophage response to TNF.

## RESULTS

### TNF triggers super-induction of IFNβ, and hyperactivity of type I interferon (IFN-I) and proteotoxic stress pathways in sst1S macrophages

Previously we have reported that B6-*sst1S* mice infected with *Chlamydia pneumoniae* (*C.p.),* as well as their bone marrow-derived macrophages (BMDMs) infected with *C.p*. in vitro, produce higher levels of IFNβ[19].This was associated with increased death of infected macrophages in vitro, which could be reduced using IFN receptor (IFNAR) blocking antibodies. To start dissecting mechanisms behind the upregulated IFNβ production, we compared IFNβ secretion by the B6wt and B6-*sst1S* BMDMs, stimulated either with a classical IFNβ inducer poly(I:C) or TNF (which induces low levels of IFNβ in B6 macrophages[24]). The B6-*sst1S* macrophages secreted higher levels of IFNβ protein in response to both stimuli (**Suppl.Fig.1A and Fig 1A**, respectively). Next, we compared the kinetics of TNF-induced IFNβ mRNA expression in B6wt vs B6-sst1S BMDMs. Initially, TNF induced similarly low levels of IFNβ mRNA expression in both cell types. Then, while IFNβ levels remained relatively stable in B6wt macrophages, in the B6-*sst1S* cells the IFNβ mRNA expression significantly increased between 8 - 24 h, such that the strain difference in IFNβ mRNA levels reached 10-20-fold by 24 h (**Fig.1B**). In addition, the B6-*sst1S* macrophages stimulated with TNF expressed significantly higher levels of the interferon-stimulated gene Rsad2 (**Suppl**. **Fig.1B)**, whose up-regulation was significantly reduced (by 70-75%) in the presence of IFNAR1 blocking antibodies, thus, confirming IFN-I pathway activation in the B6-*sst1S* cells (**Suppl.Fig.1C**). The IFNβ and Rsad2 mRNA expression kinetics demonstrated that the bias towards IFN-I pathway activation in the B6-sst1S macrophages occurred at a later stage of persistent stimulation with TNF.

**Figure 1.**
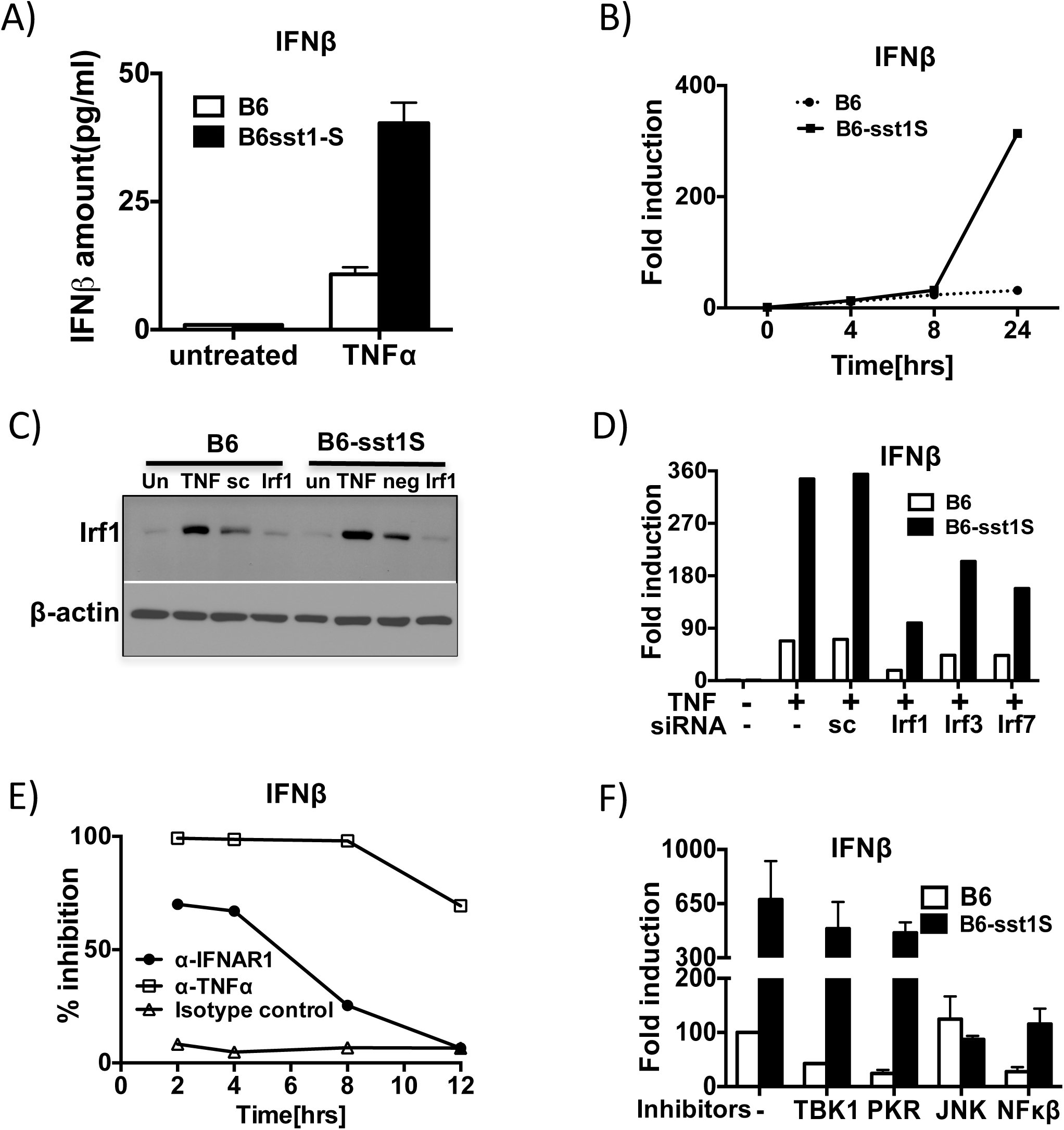
Super-induction of IFNβ in B6-sst1S BMDM after prolonged stimulation with TNF. A) IFNβ protein concentration in supernatants of B6wt and B6-sst1S BMDM treated with 10ng/ml TNFα for 24 h was detected using ELISA. Results represent data from two independent experiments. **B**) Timecourse of IFNβ mRNA expression in B6-sst1S and B6wt BMDMs after treatment with10ng/mL, as determined using real time qRT-PCR. The data are representative of three biological replicas. **C**) Effects of TNF stimulation and siRNA knockdown on IRF1 protein expression in B6 and B6-sst1S BMDMs stimulated with 10 ng/ml of TNF for 24 h. Cell were treated with siRNA 24 h prior to stimulation with TNF. Immunoblot using IRF1-specific polyclonal antibodies represents data from two independent experiments. **D**) IFNβ mRNA expression in TNF-stimulated B6 and B6-sst1S BMDM after knockdown of Irf1, Irf3 and Irf7 using siRNA (sc „ scrambled control siRNA). The data are representative of two independent biological replicas. **E**) Effect of TNF and IFN-I receptor blockade on IFNβ mRNA expression in B6-sst1S BMDM treated with 10ng/ml TNFα for 16 h. α-IFNAR1, α-TNF*α*and isotype control antibodies (10 ug/ml) were added at 2,4, 8 and 12 h of TNF treatment. IFNβ mRNA expression was calculated as % inhibition with respect to cells treated with 10ng/mL TNFα and isotype control antibodies. The data are representative of three independent experiments. **F**) Effect of small molecule inhibitors on IFNβ mRNA expression in B6-sst1S BMDM treated with 10ng/ml TNFα for 16 h. Inhibitors of TBK1, JNK, NF-κB and PKR were added after 12 h of TNF stimulation for four hours. IFNβ mRNA expression was measured by real time qRT-PCR and normalized to expression of 18S mRNA. The relative gene expression is calculated relative to the mRNA expression in untreated cells (set as 1). The data are representative of two independent experiments.

A previous report demonstrated that in wild type B6 macrophages, TNF stimulated low levels of IFNβ via NF-kB-mediated induction of IRF1, followed by auto-amplification by secreted IFNβ via the IFN-I receptor and IRF7[24]. In our model, the IRF1 protein was similarly upregulated by TNF in both B6wt and B6-sst1S mutant macrophages (**Fig.1C**). To determine which of the IRF transcription factors might play a dominant role in the IFN-I pathway hyper-activation observed in the *sst1* susceptible macrophages, we performed knockdowns of IRF1, IRF3 and IRF7 prior to stimulation of BMDMs with TNF using siRNAs (**Fig.1C** and **Suppl. Fig.1D and 1E**). IFNβ mRNA was measured at 16 h of TNF stimulation. The IRF1 knockdown had the most pronounced effect (**Fig.1D**). However, it reduced the IFNβ expression proportionally in both strains and did not eliminate the strain difference. Compared to IRF1, the IRF3 and IRF7 knockdowns had weaker effects on the IFNβ mRNA expression and also proportionally reduced the IFNβ mRNA levels in both wt and mutant macrophages (**Fig.1D**). The IFNAR1 blocking antibodies were ineffective in preventing the late phase IFNβ upregulation in the B6-sst1S cells, when added after 8 hr of TNF stimulation (**Fig.1E)**. Hence, the *sst1* locus neither acted by effects on the canonical TNF - IRF1 - IFNβ axis, nor did it control the IFNβ - IFNAR1 - IRF7 – IFNβ auto-amplification loop, described in the B6wt macrophages[24]. Although, all of those factors clearly contribute to *sst1*-independent, TNF-induced IFNβ expression in the *sst1S* macrophages as well.

To identify pathway(s) specifically responsible for the late stage super-induction of IFNβ in the B6-*sst1S* macrophages, we used small molecule kinase inhibitors. We added these agents after 12 h of TNF stimulation, and measured the IFNβ mRNA levels four hours later (**Fig.1F**). Strikingly, inhibiting JNK completely eliminated the *sst1*-dependent difference: JNK inhibitor SP600125 reduced the IFNβ mRNA expression in the B6-*sst1S* macrophages to the level of B6, but did not affect the IFNβ expression level in B6 macrophages. In contrast, NF-kB inhibitor (BAY11-7082) proportionally reduced the IFNβ mRNA levels in both wt and mutant macrophages (**Fig. 1F**). These observations suggest that the late phase super-induction of IFNβ in the *sst1*-susceptible macrophages following 12 h of TNF stimulation is a result of cooperation of the canonical *sst1*-independent TNF - NF-kB - IRF1 pathway with JNK-mediated pathway(s), presumably activated by stress, as detailed below.

To gain deeper insight into *sst1*-mediated transcriptional regulation at this critical period, we compared transcription factor (TF) activities in B6 and B6-sst1S macrophages following 12h of TNF stimulation using a TF activation array (Signosis). The activities of NF-kB, AP1, STAT1, GAS/ISRE, IRF, NFAT, NFE2, CREB, YY1 and SP1 were upregulated by TNF to a similar degree in the wt and mutant macrophages, while the HSF1 and MYC consensus sequence binding was significantly upregulated only in the mutant cells (**Fig.2A**). The higher TF activity of Myc in TNF-stimulated B6-sst1S macrophages was confirmed using EMSA (**Fig.2B**). Next, we demonstrated higher levels of c-Myc protein in the nuclei of TNF-stimulated macrophages using Western blot. Notably, the levels of nuclear c-Myc decreased between 8 and 12 hours of TNF stimulation in the B6wt macrophages, while in the *sst1*-susceptible cells nuclear c-Myc remained at elevated levels at 8, 12, and 16h after initial TNF stimulation (**Fig.2C**). Next, we tested whether increased c-Myc activity in susceptible macrophages explained increased IFNβ transcription. Indeed, knockdown of c-Myc using siRNA significantly reduced IFN*β* mRNA expression (**Fig 2D**). The small molecule inhibitor of Myc-Max dimerization 10058-F4, which suppresses E box - specific transcriptional activation by this heterodimer, i.e. targets promoter-specific TF activity of c-Myc, potently suppressed TNF-induced IFNβ and Rsad2 mRNA expression (**Fig.2E**). Myc can also promote activity of transcriptionally active genes independently of its binding to specific promoters[25, 26]. Therefore, we tested whether the inhibitor of positive transcription elongation factor (p-TEFb) flavopiridol or the RNA Pol II inhibitor of transcriptional elongation triptolide could also inhibit the IFNβ super-induction and found that both of them were inactive (**Fig.2E** and **Suppl. Fig. 1F**). In contrast, the Brd4 inhibitor JQ1, which has been previously shown to directly suppress c-Myc and IFNβ transcription[27], was active in our model as well (**Fig.2E**). In addition to transcriptional activation, c-Myc is also known to suppress a number of specific targets. Among those, cyclin-dependent kinase inhibitor p21^Cip^ plays an important role in the regulation of inflammation, monocyte differentiation, anti-oxidant response and the suppression of IFNβ gene expression[28-31]. We found that levels of p21^Cip^ were significantly reduced in susceptible macrophages during the late phase of TNF response, as compared to the wild type cells (**Fig.2F**). These data demonstrate that the susceptible phenotype is associated with persistence of c-Myc activity during response to TNF, which significantly contributes to the IFN-I pathway hyperactivation.

**Figure 2.**
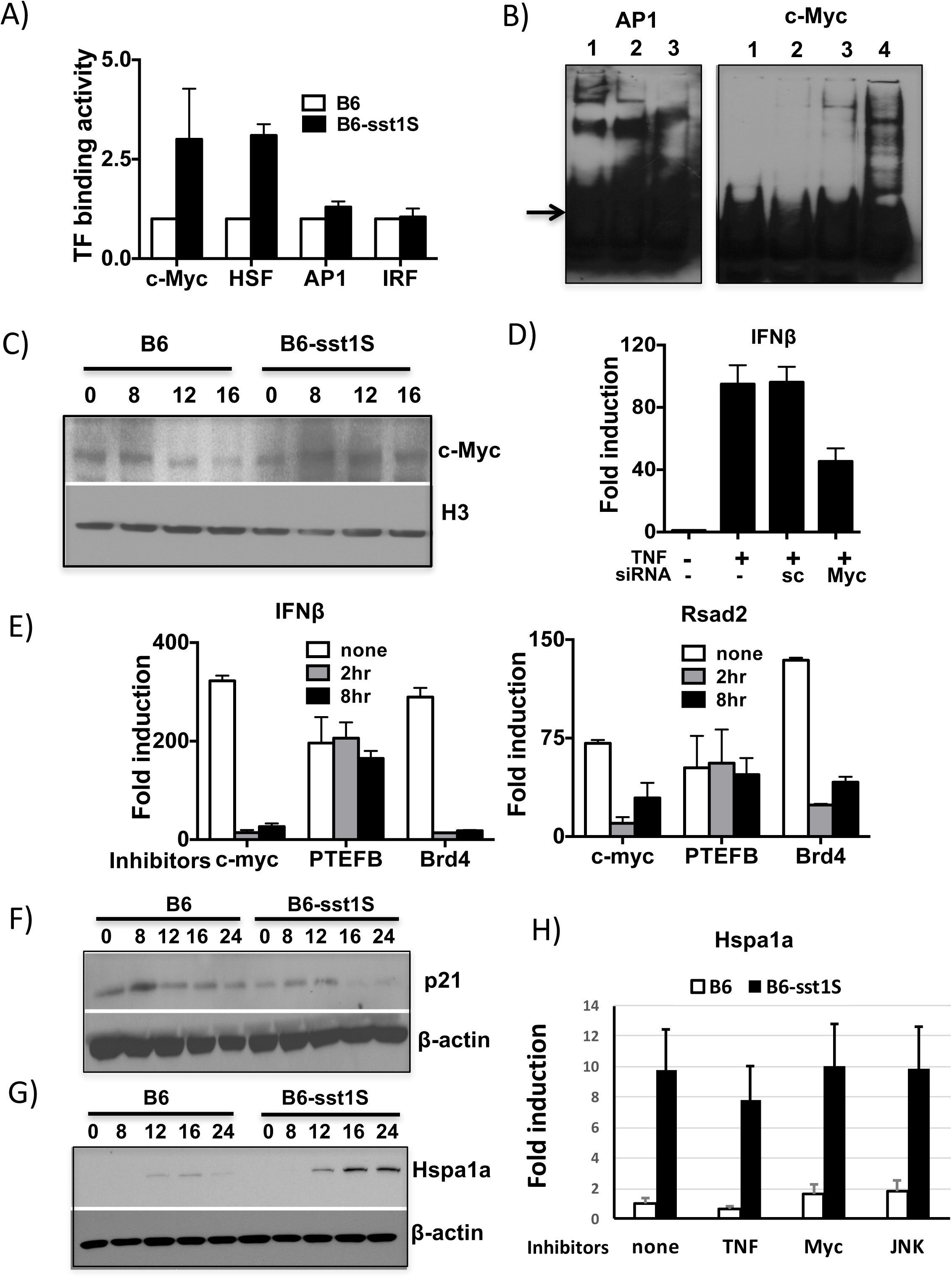
Transcriptional control of IFNβ super-induction in B6-sst1S macrophages by TNF. **A**) Transcription factor (TF) binding activities were compared using Transcription Factor Profiling Array (Signosis). Nuclear extracts (NE) were isolated from B6 and B6-sst1S BMDMs stimulated with TNF (10 ng/ml) for 12 h. Results represent data from two independent experiments. **B**) Validation of TF arrays results using EMSA with c-Myc and AP1 probes. NE were isolated at 12 h of TNF stimulation, as above. Left panel - AP-1: 1-B6wt NE, 2 „ B6-sst1S NE, 3-cold probe. The arrow denotes AP1 free probe. Right panel - c-Myc: 1-free probe, 2 „ B6wt NE, 3 „ B6-sst1S NE, 4-cold probe. Results represent data from two independent experiments. **C**) c-Myc protein levels in nuclear extracts of B6 and B6-sst1S BMDM stimulated with TNF (10ng/mL) for the indicated times (representative of two experiments). **D**). Effect c-Myc knockdown on IFNβ mRNA expression in B6-sst1S BMDM stimulated with TNF (10 ng/ml) for 18 h. **E**) Effect of inhibitors on IFNβ and Rsad2 mRNA expression in TNF stimulated B6-sst1S BMDM. Inhibitors of c-Myc (10058-F4), PTEFb (Flavopiridol) and Brd4 (JQ1) were added after 2h and 8h of TNF stimulation. IFNβ and Rsad2 mRNA expression was measured at 18 h of stimulation with TNF (10ng/mL). **F**) Timecourse of p21 protein expression in B6 and B6-sst1S BMDM treated with 10ng/mL of TNFα. The immunoblot data represent results of two independent experiments. **H)** No effects of TNF, c-Myc or JNK inhibitors added 12h of TNF stimulation on Hspa1a mRNA expression at 16 h. **G)** Time course of Hspa1a protein expression in B6 and B6-sst1S BMDMs stimulated with10ng/mL TNFα for indicated times (representative of three experiments).

In parallel, we observed escalating heat shock stress response in TNF-stimulated B6-sst1S macrophages, which is consistent with increased HSF1 activity, as determined using TF Array (**Fig.2A).** First, we confirmed the increase in HSF1 activity by demonstrating the upregulation of its transcriptional targets heat shock protein Hspa1a and Hspa1b mRNA in B6-sst1S macrophages stimulated with TNF for 24 h (**Suppl.Fig.2A**). Next, we followed the kinetics of Hspa1a protein expression and observed that initially TNF stimulation induced the heat shock stress response in both wt and mutant macrophages. However, in the wild type cells this stress response was moderate and did not escalate past 12 h, while in the susceptible cells it significantly increased between 12 and 16 h of TNF stimulation (**Fig.2G**). Taken together, the HSF1 TF activation and heat shock protein induction by TNF are indicative of proteotoxic stress (PS). The HSF1 inhibitor KRIB11, which blocks HSF1 activity, induced death of TNF-stimulated macrophages irrespective of their *sst1* genotype (**Suppl.Fig.2B**), demonstrating that, initially, PS was experienced by TNF stimulated macrophages of both backgrounds and the HSF1-mediated stress response was an important survival pathway. However, escalation of PS past 12h was characteristic of the susceptible macrophages. Of note, inhibition of TNF, c-Myc or JNK at 12 h did not prevent the PS escalation at the late phase of TNF response (**Fig.2H**), demonstrating that the incipient stress developed earlier and the PS transition to the overt phase did not require additional signaling. In contrast, the IFNβ escalation during this period still required TNF and JNK signaling (**Figs.1E and 1F**), indicating that greater PS/HSF1 pathway activation in sst1S macrophages by TNF occurred either in parallel or upstream of the IFNβ pathway. Indeed, PS is known to induce JNK activation that is a driving force behind the IFNβ super-induction seen in our model. Remarkably, another HSF1 inhibitor that, in addition to blocking HSF1, also reduces protein translation, RHT[32], not only suppressed Hspa1a mRNA expression without killing macrophages, it also eliminated the difference between the wt and sst1S macrophages, when added 2h after TNF stimulation(**Suppl.Fig.2C**). Taken together, these data demonstrate an aberrant response of the *sst1S* macrophages to TNF, characterized by imbalance of major regulators of basic cell functions, such as increased activity of c-Myc, decreased p21 protein expression and escalating proteotoxic stress, that contribute to super-induction of the type I IFN pathway.

### TNF triggers an Integrated Stress Response (ISR) and pro-apoptotic program in sst1S macrophages

#### Integrated stress response (ISR) after prolonged TNF stimulation of sst1S macrophages

To explore functional consequences of aberrant macrophage activation and more broadly characterize effects of the *sst1* locus on the late response of primary macrophages to TNF, we compared transcriptomes of B6 and B6-sst1S BMDMs after 18 h of stimulation with TNF (10 ng/ml). While no significant differences were detected in naive macrophages, the gene expression profiles of TNF-treated cells diverged substantially with 492 genes differentially expressed at p<0.001 (**Table in Fig.3A**). The most prominent differentially expressed cluster was composed of genes that were selectively upregulated by TNF in B6-sst1S, but not B6wt macrophages (**Fig.3A**). Using Gene Set Enrichment Analysis (GSEA) we found significant enrichment for the type I interferon-regulated genes in sst1S macrophages responding to TNF. Genes involved in nuclear RNA processing and nucleo-cytoplasmic transport were also upregulated by TNF in sst1S macrophages. Strikingly, multiple biosynthetic pathways were coordinately downregulated in TNF-stimulated sst1S macrophages, including lipid and cholesterol biosynthesis, protein translation, ribosome, mitochondrial function and oxidative phosphorylation. Significant upregulation of Hspa1a and Hspa1b along with IFN*β* and typical IFN-I-inducible genes Rsad2 (viperin) and Ch25h confirmed our previous observations of PS and IFN-I hyperactivity. Further validation of the differential gene expression using quantitative real time RT-PCR (qRT-PCR), demonstrated up-regulation of a number of other pathogenically important genes, such as IL-10, Mmp-13, IL-7R, Death Receptor 3 (Dr3/Tnfrsf12), transcription factors Bhlh40 and Bhlh41 (**Fig.3B**).

**Figure 3.**
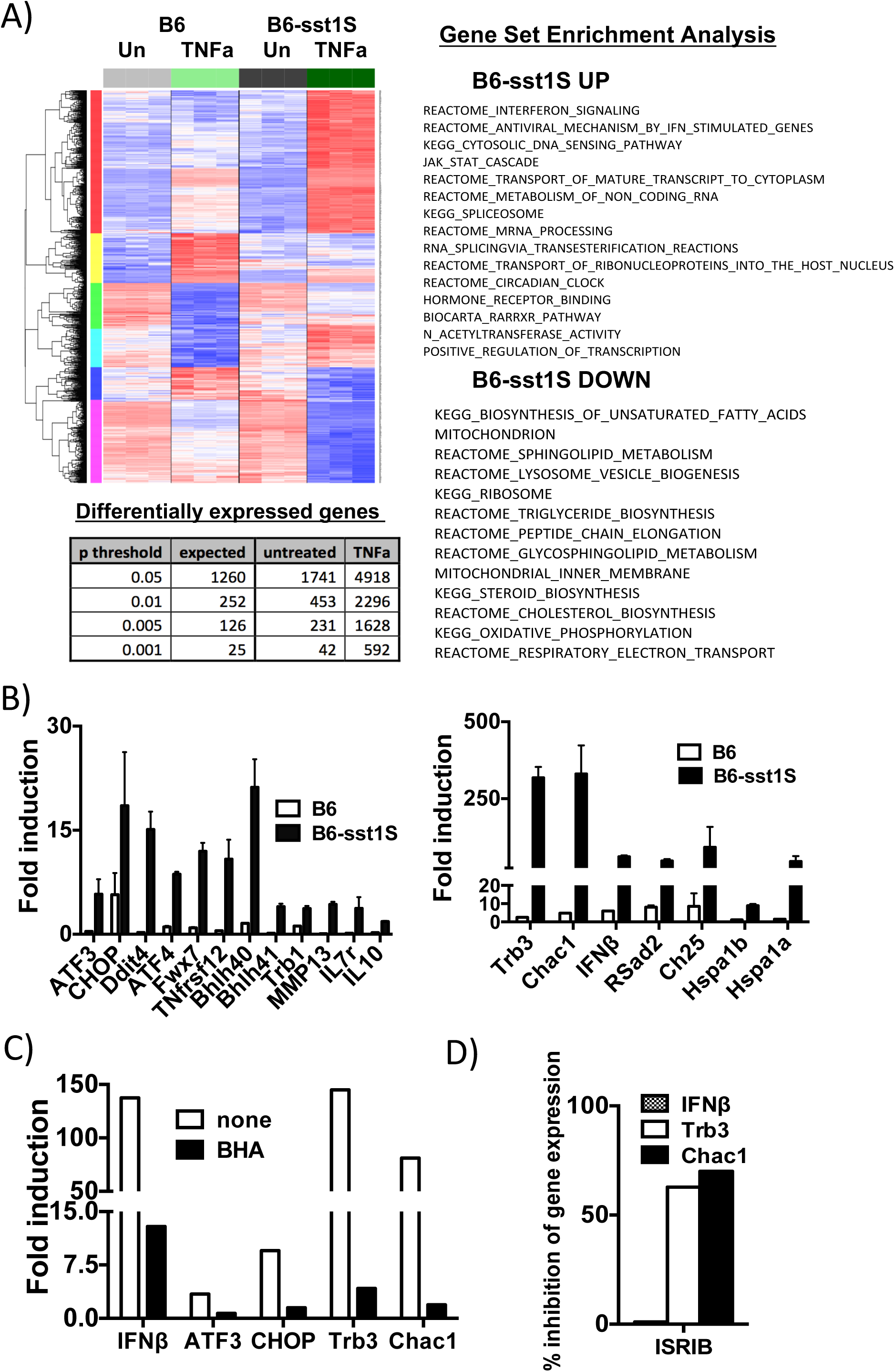
Comparison of global gene expression profiles ofB6-sst1 vs B6wt BMDMs stimulated with TNF for 18 h. **A**) Comparison of gene expression profiles of B6-sst1S vsB6 BMDM stimulated with TNF (10ng/mL) for 18 h using hierarchical clustering and gene set expression analysis (GSEA).The global gene expression was determined using Affimetrix GeneChip Mouse Gene 2.0 Arrays. **B**) Validation of microarray data using gene-specific real time qRT-PCR. **C**) Effect of ROS inhibitor BHA on IFNβ and ISR gene expression in B6-sst1S BMDM treated with 10ng/mL of TNF for 24 h. The gene expression is normalized to expression of 18S mRNA or RPS17 mRNA and presented relative to expression in untreated cells (set as 1). **D**) Effect of ISRIB on mRNA expression of IFN 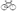 and ISR target genes Trb3 and Chac1 in B6-sst1S BMDM were treated with 10ng/mL of TNF for 24 h. Gene expression was measured by real time qRT-PCR and calculated as % inhibition with respect to cells treated with 10ng/mL TNF. The qRT-PCR results represent data from three independent experiments.

A group of genes (Atf3, Chop10, Ddit4, Trib3 and Chac1) induced during integrated stress responses (ISR) was significantly enriched among the upregulated genes (**Fig.3B**). The ISR is known to be induced as a result of the inhibition of cap-dependent translation caused by phosphorylation of eIF2*α* by several protein kinases activated in response to various stresses: viral infection (PKR), ER stress (PERK), starvation (GCN2), oxidative stress and hypoxia (HIPK)[33]. The most upregulated genes, Trib3 and Chac1, are known targets of a transcription factor Chop10 (Ddit3), which is activated downstream of the ISR transcription factors ATF4 and ATF3[34, 35]. To determine whether the induction of Trb3 and Chac1 mRNAs in the B6-sst1S macrophages by TNF was indeed downstream of eIF2*α* phosphorylation, we treated B6-sst1S BMDMs with a competitive eIF2*α* phosphorylation inhibitor ISRIB[36]. The ISRIB treatment significantly reduced the Trb3 and Chac1 mRNA upregulation, thus confirming their specific induction by ISR in our model. However, ISRIB had no effect on IFNβ mRNA level (**Fig.3D**). In contrast, addition of the ROS scavenger BHA immediately after TNF stimulation inhibited both ISR and IFNβ gene expression, indicating that both pathways developed in response to oxidative stress (**Fig.3C).** Indeed, ROS are known to cause protein misfolding and aggregation in cytoplasm and ER. The levels of ROS produced by TNF-stimulated B6wt and B6-sst1S macrophages were similar (**Suppl.Fig.2D**) suggesting that those cells may differ in responses to ROS-mediated stress. Hence, we used Trb3 and Chac1 mRNA expression as biomarkers of ISR to interrogate mechanisms of the ISR activation and maintenance in sst1S macrophages during the course of TNF stimulation.

#### Bi-phasic regulation of the Integrated Stress Response

First, we compared the mRNA kinetics of genes representing transcriptional targets of the ISR (Chop10, Atf3, Ddit4, Chac1 and Trb3). The expression of the ISR genes spiked in the B6-sst1S cells at 16 h and continued to increase further between 16 - 24 h (**Fig.4A**). Next, we monitored the expression of ISR markers ATF4, ATF3 and GADD34 at the protein level by Western blot. Initially, we observed similar induction of ATF4 and ATF3 after 3 h of TNF stimulation in both the B6wt and B6-sst1S BMDMs. However, in the B6 cells the levels of ATF4 and ATF3 proteins declined to basal levels by 15 and 24 h, respectively. Meanwhile, in the susceptible macrophages they remained elevated. The ATF3 levels even increased during the 16 - 24 h interval (**Fig.4B**). Thus, the sst1 susceptibility allele is associated with ISR escalation after 12hrs of TNF stimulation, i.e. resembles its effect on IFN-I pathway suggesting a mechanistic link.

We followed the kinetics of the ISR-and IFN-inducible genes within a critical period between 8 and 14 hours at 2 h intervals. The IFNβ mRNA expression level in the B6-sst1S macrophages gradually increased, while the ISR markers remained at the same level throughout this period, suggesting a possible mechanistic hierarchy (**Fig.4C**). Therefore, we tested whether blocking IFN-I signaling reduced the ISR induction. The IFN type I receptor (IFNAR1) blocking antibodies were added at different times after stimulation with TNF (10 ng/ml), and the ISR was assessed at 16 h of the TNF stimulation (**Fig.4D**). Blocking the IFN-I signaling at 2 - 4 h after TNF stimulation profoundly suppressed the ISR escalation, as measured by Trb3 and Chac1 gene expression at 16 h of TNF stimulation. However, the effect of IFNAR blockade at later timepoints gradually declined and completely disappeared by 12 h. Blocking TNF signaling using neutralizing antibodies affected the ISR induction in a manner very similar to the IFNAR blockade - it was effective at the earlier stages (2-4 h) and only partially efficient at 8 h (**Fig.4D**). Simultaneous blockade of both pathways at 8 h did not produce synergistic effect on ISR suggesting that both TNF and IFN were in the same pathway (**Suppl.Fig.3A**). The effect of ROS scavenger on ISR escalation dramatically declined by 8 h. Effects of the above inhibitors on ISR completely disappeared by 12 h (**Fig. 4D and Suppl.Fig.3B**). Therefore, once the TNF - IFN- and ROS-dependent ISR pathway was set in motion, the transition from latent to overt ISR during the 12 - 16 h period was driven in cell autonomous manner.

**Figure 4.**
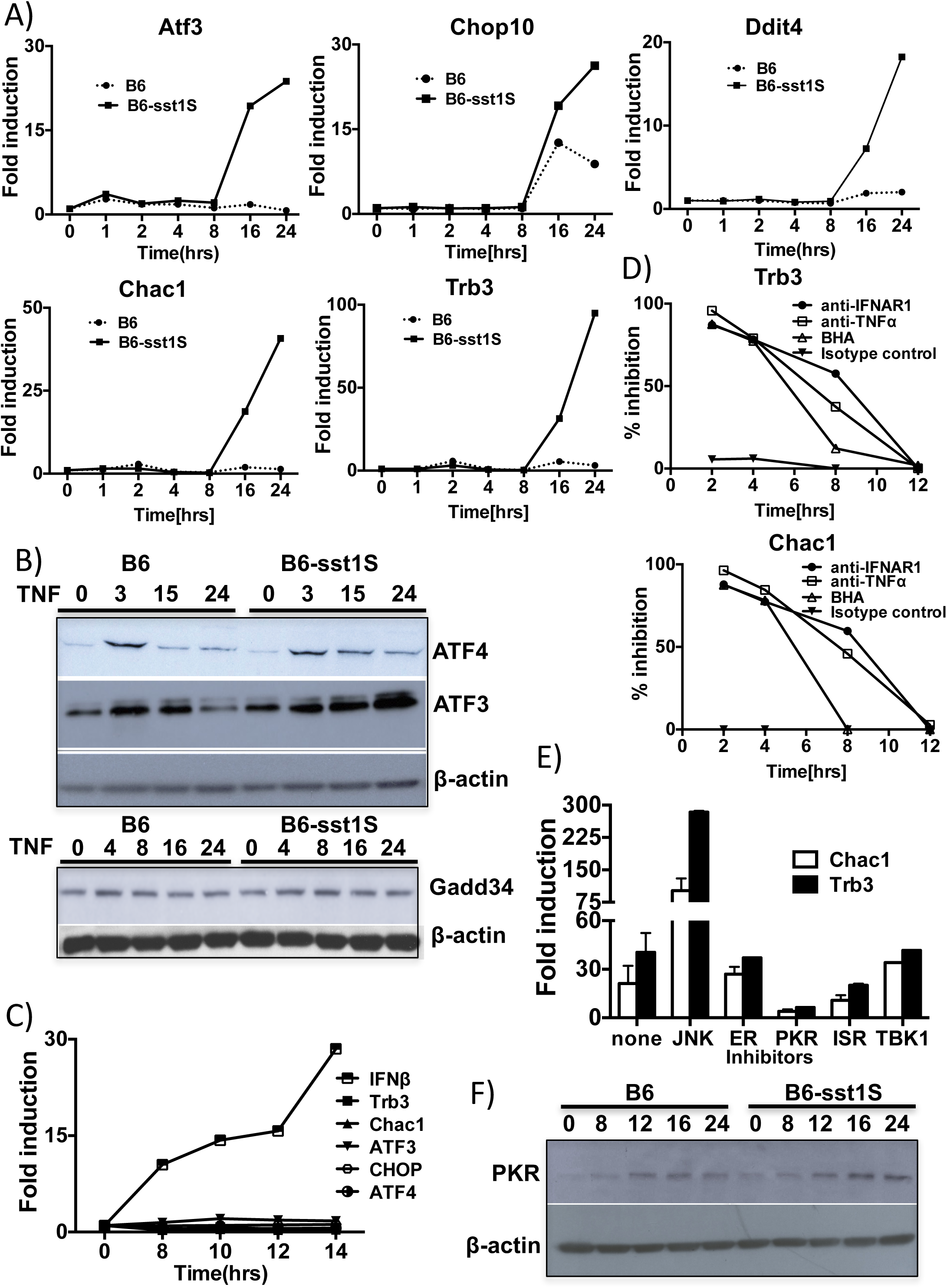
TNF treatment leads to bi-phasic upregulation of integrated stress response in B6-sst1S BMDM. **A**) Timecourses of the ISR gene mRNA expression: Atf3, Chop10, Chac1, Trb3 and Ddit4 mRNA expression levels in B6-sst1S and B6wt BMDMs after treatment with TNF (10ng/mL). **B**) Timecourse of ISR protein expression in TNF-stimulated macrophages. Immunoblot analysis of ATF3, ATF4 and GADD34 levels in whole cell extracts of B6wt and B6-sst1S BMDM treated with 10ng/mL of TNFα for the indicated times. **C**) Real time qRT-PCR analysis of the mRNA kinetics of IFNβ and ISR genes in B6-sst1S BMDMs after stimulation with10ng/mL TNF for 8, 10, 12 and 14 h. Real time PCR data is normalized to expression of 18S mRNA mRNA and presented relative to expression in untreated cells (set as 1). **D**) Time dependent effects of TNF and IFNAR blockade and ROS inhibition on Trb3 and Chac1 mRNA expression in B6-sst1S BMDM treated with 10ng/ml TNFα for 16hrs. α-IFNAR1, α-TNFα, BHA and isotype control antibodies were added after 2, 4, 8 and 12 h of TNF treatment. Trb3 and Chac1 mRNA expression was calculated as % inhibition with respect to cells treated with 10ng/mL TNFα. **E**) Effect of inhibitors on late phase ISR gene expression in TNF stimulated B6-sst1S BMDM. Inhibitors of JNK, ER stress (PBA), PKR, ISR and TBK1 were added after 12h of TNF stimulation (10 ng/mL), and Trb3 and Chac1 mRNA levels were measured at 16h. Trb3 and Chac1 mRNA expression was normalized to expression of 18S rRNA and presented relative to expression in untreated cells (set as 1). The qPCR results represent data from three independent experiments. **F**) Timecourse of PKR protein expression in B6-sst1S and B6wt BMDMs treated with TNF (10ng/mL) for indicated times. The Western blot is representative of two independent experiments.

To reveal the driving force behind the ISR transition from the latent to overt phase, we treated B6-sst1S BMDM at 12 h after stimulation with TNF with inhibitors of eIF2*α* phosphorylation ISRIB, ER stress inhibitor PBA[37], PKR inhibitor C16[38] and inhibitors of stress kinases p38 (SB203580) and JNK (SP600125) and measured the induction of Trb3 and Chac1 mRNAs at 16 h (**Fig.4E**). The ISRIB and PKR inhibitors profoundly inhibited expression of both sentinel mRNAs, while PBA had no effect suggesting that PKR activity was responsible for transition from latent to overt ISR in the B6-sst1S macrophages at the late stage. Indeed, PKR levels increased by the 12 h of TNF stimulation and were subsequently maintained at higher level in the B6-sst1S cells (**Fig.4F**). PKR is a classical interferon-inducible protein, whose kinase activity is induced by double-stranded RNA (dsRNA). Recently it has been demonstrated that PKR can interact with and be activated by small nucleolar RNAs and other misfolded and dimerized endogenous RNA molecules [39, 40]. Using dsRNA-specific antibodies J2[41], we detected dsRNA speckles in the cytoplasm of TNF-stimulated BMDM of both B6 and B6-sst1S backgrounds (**Suppl. Fig.3C**). Thus, the presence of the endogenous PKR ligands may provide an explanation of how IFN-induced PKR is activated by TNF in non-infected macrophages.

Taken together, these data demonstrate that in sst1S macrophages ROS induced by TNF initiated a cascade of stress responses in biphasic manner. The early initiation phase (2 „ 4 hrs) required ROS, leading to proteotoxic stress (PS), ER stress and ER stress-mediated ISR similarly in both the wt and sst1S mutant macrophages. In support of this notion, XBP-1 splicing, known to be induced by Ire1 kinase activated specifically by the ER stress follows similar kinetics in both strains reaching peaks at 4 - 8 h (**Suppl. Fig.3D**). While self-limited in the B6wt cells, the ISR escalated in the B6-sst1S macrophages during the 12 - 16 h period of TNF stimulation in IFN-I-dependent manner via PKR activation, a pathway traditionally associated with antiviral immunity, but more recently also linked to metabolic dysregulation[42].

#### TNF induces pro-apoptotic reprograming in the sst1S macrophages

The major adaptive role of ISR is a global reduction of cap-dependent protein translation[34]. However, translation of many proteins involved in stress responses proceeds via cap-independent mechanisms and the proportion of those proteins in the cellular proteome increases during prolonged stress. Thus, we postulated that an unrelenting stress in B6-sst1S macrophages would result in global proteome remodeling, and discordance between transcriptome and proteome. Indeed, we detected upregulation of Trb3 protein at the early stage of TNF stimulation, but not during the second phase of ISR escalation in B6-sst1S macrophages, despite high levels of its mRNA induction (**Fig.5A**). Therefore, we compared global quantitative protein abundance profiles of the B6wt and B6-sst1S mutant macrophages after stimulation with TNF, using stable isotope labeling of the digested macrophage proteomes with tandem mass tags (TMT) followed by deep 2-D LC-MS/MS-based proteomic analysis.

**Figure 5.**
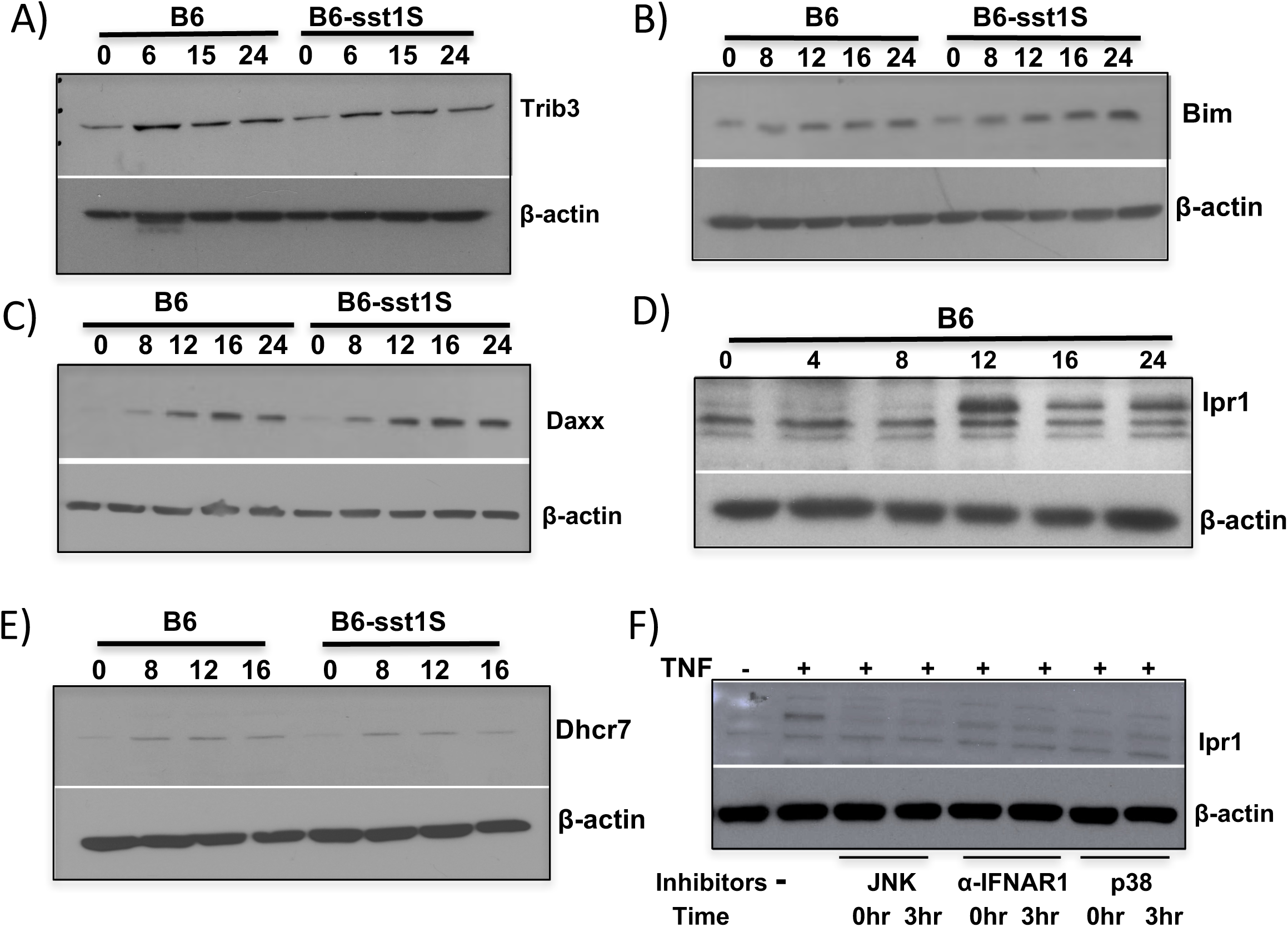
Proteins differentially expressed in TNF-stimulated B6-sst1S and B6wt macrophages. Validation of proteomics data using immunoblot analysis and the kinetics of Trib3(**A**), Bim(**B**), Daxx(**C**), Ipr1(**D**) and DHCR7(**E**) proteins in B6-sst1S and B6wt BMDMs treated with10ng/mL of TNF. Whole cell extracts were isolated at indicated times. **F**) Effect of JNK and p38 inhibitors and IFNAR blockade on IPR1 protein induction in B6wt BMDM treated with 10ng/mL of TNFα for 24 h. The inhibitors were added at the beginning (0hr) or after 3hrs of TNF stimulation. Immunoblotting was carried out using Ipr1 polyclonal antibody and represents two independent experiments.

The proteome profiles of TNF-stimulated B6-sst1S and B6wt macrophages were clearly distinct. A number of proteins were up-regulated in the susceptible macrophages demonstrating the absence of a total translational arrest in the mutant cells. In agreement with our previous observations, higher levels of ATF3 and Hspa1a proteins were detected by the proteomic analysis. We also observed that the TNF-stimulated mutant sst1S cells expressed higher levels of proapototic proteins DAXX and Bim, cold shock-inducible RNA binding protein Rbm3 and dsRNA-binding protein Stauphen1. We confirmed the upregulation of DAXX and Bim by Western blot (**Fig.5B and C**).

Only the B6wt cells expressed the Sp110/Ipr1 protein encoded within the sst1 locus. Ipr1 has been identified and validated as a strong candidate gene in our previous work using positional cloning[20]. The Ipr1 protein was present in proteome of both non-stimulated and, at higher levels, of TNF-stimulated B6 macrophages. We extended these observations by finding that the Ipr1 protein was induced in B6wt macrophages between 8 and 12 h after initial TNF stimulation, corresponding to a period of late stress escalation in the Ipr1-negative B6-sst1S cells (**Fig.5D**). JNK inhibition or IFNAR blockade prevented the Ipr1 protein up-regulation by TNF demonstrating that it is an interferon- and stress-inducible protein (**Fig.5F**). The TNF-stimulated B6 macrophages also expressed higher levels of proteins involved in antioxidant defenses and protein homeostasis in ER and cytoplasm: 1) NADH-cytochrome b5 reductase 4 (CYB5R4) which protects cells from excess buildup of ROS and oxidant stress[43]; 2) Stromal cell-derived factor 2 (SDF2), involved in ER protein quality control, unfolded protein response and cell survival under ER stress[44]; 3) The signal sequence receptor 2 (SSR2), a subunit of the ER TRAP complex involved in protein translocation across the ER membrane[45]; 4) Stress-associated endoplasmic reticulum protein 1 (SERP1) which interacts with target proteins during their translocation into the lumen of the endoplasmic reticulum and protects unfolded target proteins against degradation during ER stress[46]. In addition, the B6 macrophages expressed higher levels of 7-dehydrocholesterol reductase (Dhcr7), a key enzyme in cholesterol biosynthesis. We confirmed this observation using Western blot and determined that the difference was due to the down-regulation of Dhcr7 in the B6-sst1S macrophages 12 - 16 h of TNF stimulation (**Fig.5E**). This effect might be explained by the inhibitory effect of 25-hydoxycholesterol on cholesterol biosynthesis via SREBP2 inhibition[47]. This oxidized cholesterol derivative is produced by the IFN-I -inducible enzyme Ch25h (cholesterol 25-hydroxylase)[48], which is highly upregulated in the B6-sst1S macrophages by TNF in an IFN-I-dependent, but ISR-independent manner. Thus, hyperactivity of IFN-I pathway in the sst1-susceptible macrophages leads to pro-apoptotic proteome remodeling via up regulation of PKR-ISR-mediated pathway, and, possibly, broader down regulation of metabolic pathways, such as cholesterol biosynthesis, via effects of other interferon-stimulated proteins. In contrast, the proteome of B6wt macrophages stimulated with TNF is enriched for proteins countering oxidative and ER stress and supporting survival.

### TNF priming modifies susceptibility of the sst1S macrophages to infections in vitro

In vivo, monocytes are recruited to and undergo terminal differentiation within inflammatory milieu, where they will encounter cytokines prior to interaction with infectious agents. The above data suggested that pre-exposure of macrophages to TNF may profoundly affect their subsequent interactions with various pathogens, not only Mtb. Therefore, we tested whether pre-treatment of B6 and B6-sst1S macrophages with TNF differentially affected their interactions with another intracellular bacterial pathogen, *Francicella tularensis* Live Vaccine Strain (F.t. LVS). Following overnight pretreatment with10 ng/ml of TNF, the death rate of the infected B6-sst1 macrophages increased at various MOIs (**Fig.6A**). It was significantly higher in the B6-sst1 background, as compared to B6wt (**Fig.6B**). The bacterial loads also increased in TNF pre-treated susceptible macrophages (**Fig.6C**) suggesting that stress compromised their anti-bacterial resistance. Assessing the levels of IFNβ, Rsad2 and stress mRNAs in macrophages, either pretreated with TNF or naïve, we determined that TNF treatment made a major contribution to IFN and stress pathways induction, as compared to the bacteria alone (**Fig.6D**).

**Figure 6.**
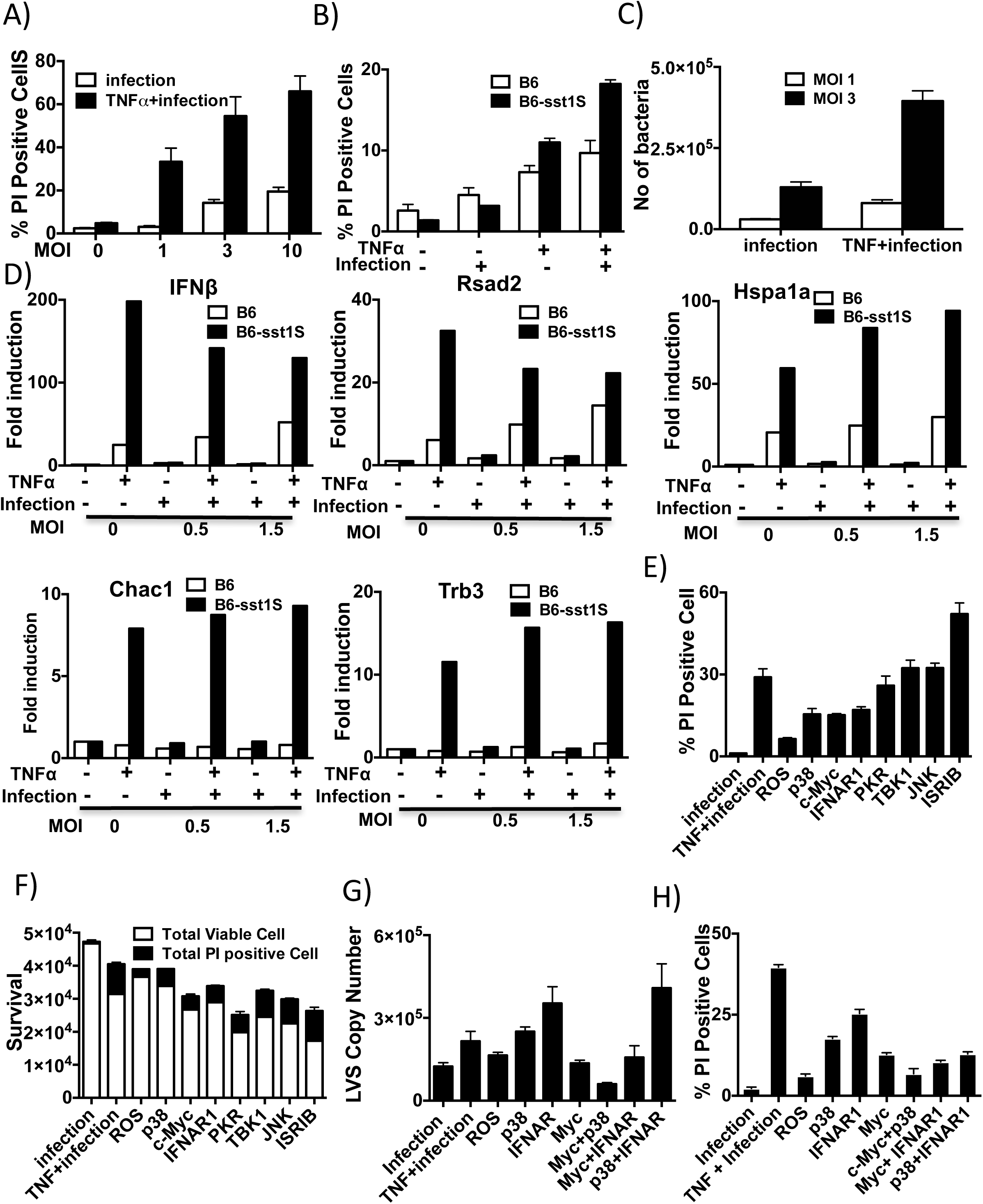
Effect of TNF priming on infection with *F.t.* LVS in sst1S BMDM. **A)** Survival of B6**-**sst1S BMDM, either naïve (control) or primed with 10ng/mL of TNF*α* for 16 h, after infection with *F.t.* LVS at MOI 1, 3 and 10 for 24 h. Percentage of PI-positive cells was determined using automated microscopy (Celigo). **B**) Effect of the sst1 locus on survival of B6-sst1S and B6wt BMDM treated with TNF and infected with *F.t.* LVS at MOI =1. Cell death was measured using % of PI positive cells, as above. **C)** Effect of TNF priming on *F.t.* LVS control by the B6**-**sst1S BMDM. The macrophages, either naïve or primed with 10ng/mL of TNF*α* for 16hrs, were infected with *F.t*. LVS at MOI 1 and 3 for 24 h. The bacterial loads were determined using qPCR. **D**) Comparison of the effects of TNF and *F.t.*LVS infection on IFN-I and stress response gene expression in B6-sst1S BMDM. Macrophages, either naïve or primed with 10ng/mL of TNFα for 16 h, were infected with *F.t.* LVS at MOI 0.5 and 1.5 for 24hrs. IFNβ, Rsad2, Hspa1a, Trb3 and Chac1 mRNA expression was measured using qRT-PCR. Real time PCR data is normalized to expression of 18S RNA and presented relative to expression in untreated cells (set as 1). **E**) Effect of small molecule inhibitors on cell death of B6-sst1S BMDM primed with 10ng/mL of TNFα for 16 h and infected with *F.t.* LVS at MOI=1 for 24hrs. The inhibitors were added after 4hrs of TNF treatment. Cell death was measured as % of PI positive cells using automated microscopy. **F)** Total (live and dead) numbers of cells treated as in E) was estimated by measuring total number of viable and dead (PI+) cells. **G-H)** Effects of the pathway inhibitors and their pairwise combinations on bacterial loads (**G**) and macrophage cell death (**H**) in TNF-stimulated B6-sst1S BMDM infected with *F.t.*LVS (MOI=1). Cells were primed with TNF, infected with *F.t.* LVS and treated with inhibitors as described above. All cell death data is representative of three independent experiments. Bacterial enumeration was done using real time qPCR and was normalized to total cell number determined using automated microscopy. All qPCR results represent data from three independent experiments.

To assess the impact of individual stress pathways on the LVS-infected macrophage survival and bacterial control, we pre-treated B6-sst1S BMDMs with TNF in the presence of small molecule inhibitors and then infected them with *F.t.* LVS at MOI 1:1 and 3:1. In this experiment, TNF and inhibitors were present for the duration of infection (24h), as opposed to short stage-specific pulses of inhibitors used in previous experiments. This modification was introduced to allow detection of integrated effects of inhibitors on cell survival and bacterial control. First, we determined that inhibitors of ROS (BHA), p38, c-Myc, as well as IFNAR blocking antibodies, significantly reduced percentage of PI positive cells (**Fig.6E**) and prevented cell death (**Fig.6F**). However, the ISR inhibitor ISRIB and, to a lesser extent JNK and PKR inhibitors aggravated the susceptible phenotype, as evidenced by significant decrease of total and viable cell numbers after infection (**Fig.6F**). This observation points to compensatory roles of JNK and ISR stress pathways activation in the susceptible background, i.e. they function to improve cell survival. Therefore, those inhibitors were excluded from subsequent experiments, in which we studied effects of inhibitors on the bacterial control. As above, B6-sst1S BMDMs were treated with TNF in the presence of individual ROS, p38, c-Myc inhibitors and IFNAR blocking antibodies, as well as their pairwise combinations. Remarkably, the c-Myc inhibitor significantly reduced, while the IFNAR1 receptor blockade increased, the bacterial loads in TNF treated B6-sst1S macrophages, while ROS and p38 inhibitors were neutral in this respect (**Fig. 6G**). In pairwise combinations, c-Myc and p38 inhibitors improved the bacterial control and increased macrophage survival (**Figs.6G and 6H**, respectively). Also, the c-Myc, but not the p38, inhibitor could overcome detrimental effect of the IFNAR blockade on the bacterial replication. Overall, our data demonstrate that pre-exposure of B6-sst1S macrophages to TNF prior to infection with virulent intracellular bacteria (*F.t.* LVS is virulent in mice) in vitro, compromises infected macrophage survival and bacterial control via the same mechanisms that underlie TNF-induced stress: in agreement with our previous data demonstrating that Myc hyperactivity mechanistically is upstream of other manifestations of aberrant activation in B6-sst1S macrophages, inhibition of Myc transcription factor activity uniquely improved both macrophage survival and bacterial control.

### The *sst1* locus regulates outcome of infection with *F. tularensis* LVS in vivo

To assess the impact of the *sst1* locus in vivo, we infected the *sst1* resistant and susceptible congenic mice via respiratory route with 1000 CFU of *F.t.* LVS. The survival of the *sst1* susceptible mice was significantly lower as compared to the *sst1*-resistant congenic mice (**Fig. 7A**). Importantly, the bacterial replication was initially similar in the lungs, spleens and livers of both mouse strains. The bacterial loads, however, significantly diverged between days 5 and 11 reaching 100-fold higher levels in the organs of the *sst1* susceptible mice (**Fig.7B and Suppl. Fig. 4A**), which also developed extensive necrotic lung inflammation by that time (**Fig.7C and Suppl. Fig 4B**). Additional experiments were performed using *sst1* congenic *scid* mice also developed in our laboratory. Both the sst1R *scid* and sst1S *scid* mice succumbed to infections with 60 CFU of *F.t.* LVS. However, the survival time was shorter and the bacterial loads were 30-50-fold higher in the *sst1* susceptible *scid* mice (**Suppl. Fig.5A and 5B**) demonstrating that innate immune cells were responsible for the *sst1*-mediated phenotype *in vivo.*

**Figure 7.**
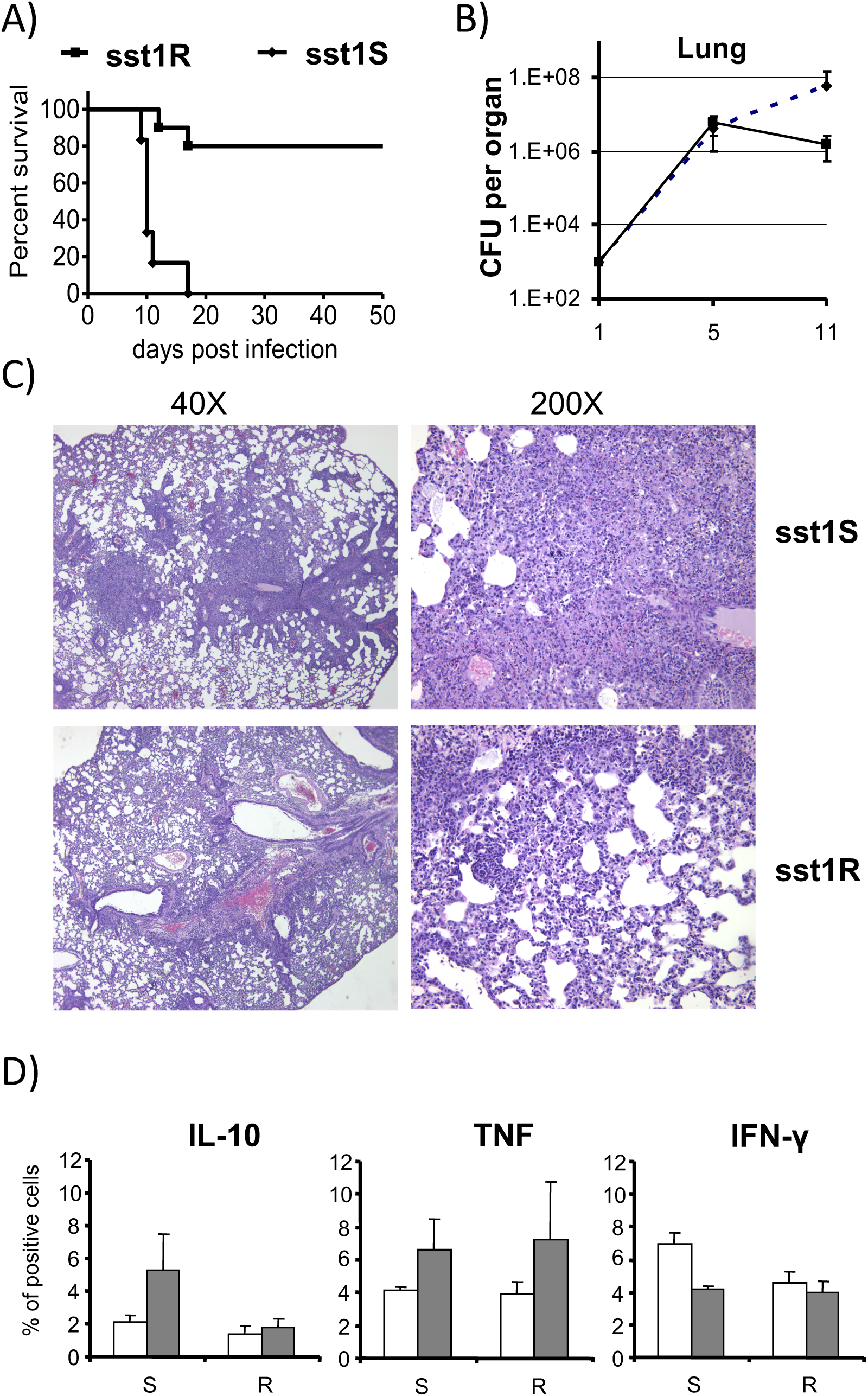
The sst1 locus controls host resistance to aerosol infection with *F.t.* LVS. **A**) Survival of the *sst1^R^* and *sst1^S^* inbred mouse strains after aerosol infection with 1600 CFU of *F.t.* LVS. Six mice per group were used in each strain. **B**) Kinetics of *F.t.* LVS growth in the lungs of the *sst1^R^* and *sst1^S^* mice after the aerosol infection with 1,600 CFU of F. LVS. Four mice per group were sacrificed at each time point for CFU determination using plating of serial dilutions of lung homogenates. **C**) Histopathology of the lungs of *sst1^S^* (upper panels) and *sst1^R^* (lower panels) 11 days post aerosol infection with 1,600 CFU of F. LVS. H&E staining, magnification X40 (left panels) and X200 (right panels). **D**) Intracellular cytokine staining of the lung cells isolated 5 (white bars) or 10 (grey bars) days post aerosol infection with 300 CFU of *F.t.* LVS. Three mice per group were used in the experiment. S - sst1^S^; R - sst1^R^. Data is representative of two independent experiments.

FACS analysis of inflammatory cells isolated from the infected lungs of immune competent mice eight days post infection demonstrated similar proportions of CD4^+^ and CD8^+^ T cell populations and NK cells. The proportions of activated CD69+ cells within those populations were also similar (**Table 1**). However, we detected a higher proportion of the IL-10 producing myeloid cells in the *sst1S* mouse lungs at that time, which is in agreement with the BMDM phenotype observed in vitro (**Fig.7D**). In addition, we observed substantial difference in the myeloid compartment (CD11b+), where the fraction of immature monocyte-like Ly6C^+^F4/80^−^ cells was significantly higher in the lungs of the *sst1S* mice (11.2%), as compared to 5.3% in the B6 lungs. The ratio of the more mature (Ly6C^+^F4/80^+^) monocyte-derived macrophages to the immature (Ly6C^+^F4/80^−^) cells in the lungs of the *sst1*S mice was 1.1, as compared to 3.7 in the resistant ones (**Table 2)**. These data imply either delayed maturation of the newly recruited inflammatory monocytes in the lungs of the *sst1S* mice and/or more rapid turnover of monocyte-derived macrophages during *F.t.* LVS infection *in vivo*, which would be consistent with their more rapid demise, as observed in vitro.

**Table 1:**
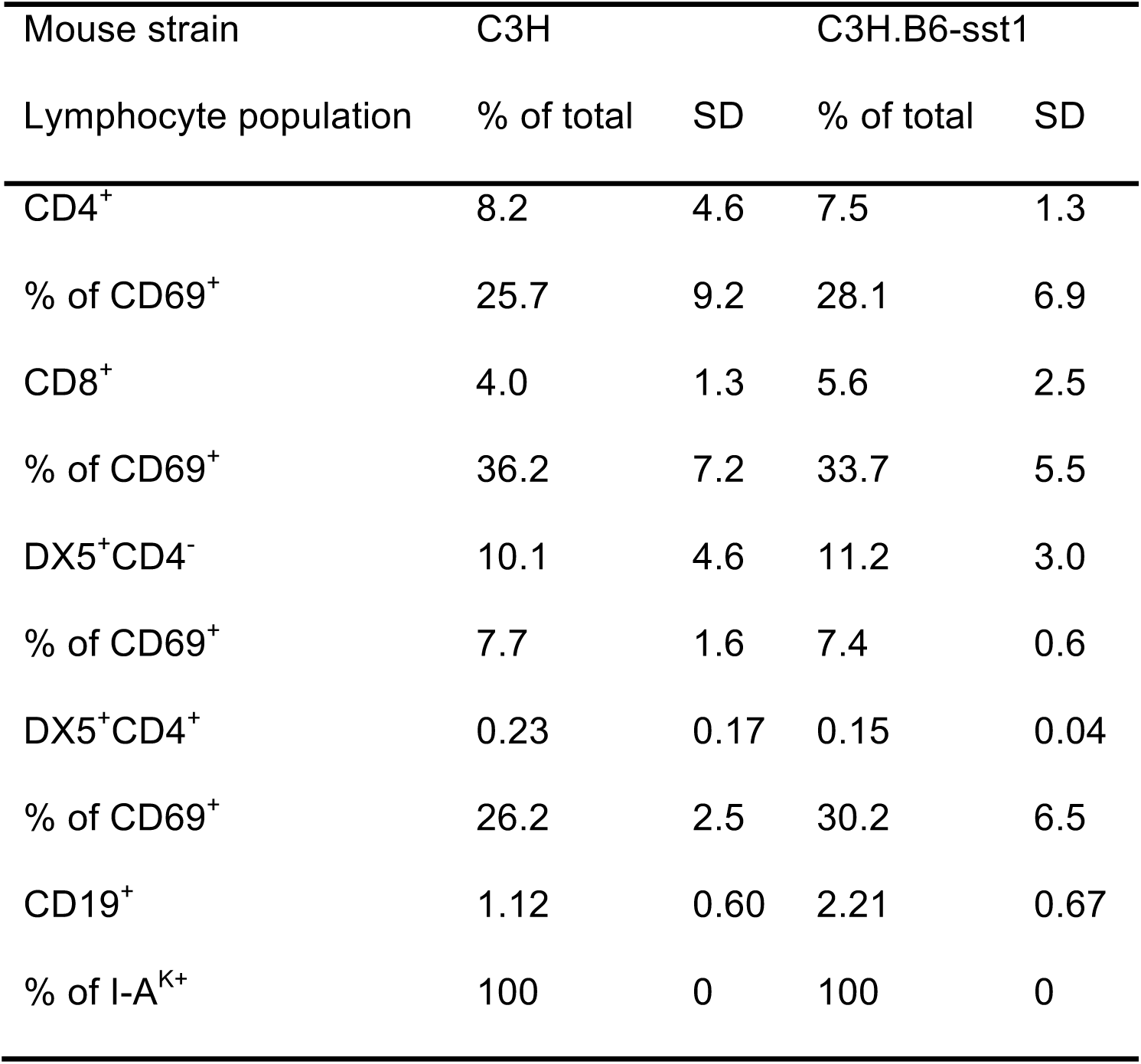
Flow cytometry of lymphoid lung cells 8 days following aerosol challenge of C3H and C3H.B6-sst1 mice with 300 CFU of *F.t.* LVS

**Table 2:** Flow cytometry of myeloid lung cells 8 days following aerosol challenge of C3H and C3H.B6-sst1 mice with 300 CFU of *F.t.*LVS

| Mouse strain | C3H |  | C3H.B6-sst1 |  |
| --- | --- | --- | --- | --- |
| Myeloid cell population | % of total | SD | % of total | SD |
| F4/80<sup>-</sup>Ly6C<sup>+</sup> | 11.2 | 2.0 | 5.3 | 3.7 |
| F4/80<sup>+</sup>Ly6C<sup>+</sup> | 12.7 | 4.4 | 19.5 | 7.2 |
| F4/80<sup>+</sup>Ly6C<sup>-</sup> | 0 | 0 | 0 | 0 |
| Ly6G<sup>+</sup>F4/80<sup>-</sup>Ly6C<sup>-</sup> | 39.3 | 9.5 | 37.7 | 5.2 |

## DISCUSSION

Taken together, the complex cascade delineated in this study indicates that the *sst1*-mediated susceptibility to intracellular bacterial pathogens is mechanistically linked to an aberrant macrophage response to TNF triggering and sustaining escalating stress responses. IFN type I plays an important intermediary role in this cascade, with its super-induction resulting from a synergistic effect of the canonical NF-kB-mediated TNF activation pathway and a *sst1*-specific response mediated by the stress kinase JNK. The origins of stress, however, are upstream of IFN-I and reflect hyperactivity of Myc in the setting of TNF-induced oxidative stress. Inhibition of either transcriptional activity of Myc or ROS generation prevented the late phase IFN-I super-induction and the escalation of the ISR in TNF-stimulated sst1S macrophages. Thus, we suggest that untangling stress-mediated regulation of Myc activity in macrophages is a key to understanding the *sst1*-mediated phenotype in vitro and in vivo.

The *sst1* locus encodes interferon-inducible nuclear protein Sp110, whose expression is completely abolished in TNF-or IFN*γ*-activated macrophages carrying the *sst1* susceptible allele[20]. In the wild type macrophages, this protein is induced at the late stage of TNF response between 8 – 12 h. This time interval is critical for the expression of the susceptible phenotype, because it precedes the escalation of the proteotoxic stress and super-induction of IFNβ. During this period, Sp110 accumulates in the nucleus and associates with chromatin (Bhattacharya, unpublished). We hypothesize that this protein may play direct or indirect roles in limiting Myc activity in classically activated macrophages. Conversely, the *sst1* congenic interval encodes additional members of the Sp100 family (Sp100 and Sp140), as well as other genes expressed in macrophages that may participate in regulating multifaceted Myc activities. To our best knowledge, none of the *sst1*-encoded gene products have been implicated in cross talk of stress-, interferon-and Myc-mediated pathways, so far. Thus, the molecular mechanisms that negatively regulate Myc-driven programs in activated macrophages and controlled by genes encoded within the *sst1* locus remain to be elucidated.

While Myc activity is important for growth of myeloid precursors[49] and alternative macrophage activation in tumor microenvironment[50], down regulation of Myc activity in monocytes/macrophages appears to be an important adaptive mechanism within infection-induced inflammatory lesions. The inflammatory monocytes are produced from myeloid precursors in bone marrow and recruited from circulation to sites of inflammation, where they undergo final cell divisions and terminal differentiation. Prior to pathogen encounter, their phenotype is further sculpted by multiple factors within the inflammatory milieu, which include not only classic inflammatory mediators, but also growth factors, hypoxia, starvation, acidosis, etc. Sensing and integrating responses to gradients of those factors is necessary to balance cell metabolism, growth and macrophage effector functions within specific environments[11]. Indeed, in non-transformed cells, sensing stress can stop cell cycle progression and trigger terminal differentiation[51]. For example, stress kinases can stimulate p53 to block transcription of c-Myc and its targets[52]. Considering macrophage maturation within inflammatory lesions, this response may dramatically reduce energy expenditure and prepare macrophages for subsequent stress escalation and pathogen encounter. Indeed, we determined that Myc nuclear levels and transcriptional activity was down regulated 12 hours after TNF stimulation, but only in the wild type macrophages. In contrast, the inability to repress Myc in TNF-stimulated susceptible macrophages is associated with escalation of proteotoxic stress (PS), possibly due to misfolding of newly synthesized proteins caused by ROS. Thus, persistent transcriptional activity of Myc appears to be a major driver of the maladaptive response of the sst1-susceptible macrophages, leading to un-resolving stress, accelerated death of infected macrophages and, eventually, to the development of overt necrotic pathology.

Our studies revealed the dynamics of downstream cascade associated with the susceptible phenotype. At an early stage, TNF stimulation causes proteotoxic stress (PS) and integrated stress response (ISR) in both wild type and the *sst1*-susceptible mutant macrophages. At this stage the ISR is driven by the ER stress and unfolded protein response (UPR), as previously described[53]. The unusual, second wave of ISR activation is exclusive for the *sst1*-susceptible phenotype. It is initiated by TNF in IFN-I-dependent manner via PKR activation, similar to a cascade recently described in a model of *Listeria monocytogenes* infection[54]. However, in the *sst1*-susceptible macrophages, the IFN*β* super-induction was triggered by TNF alone. We excluded a significant contribution of the STING – TBK1 - IRF3 pathway to the observed IFN*β* super-induction and, thus, demonstrated that recognition of endogenous or exogenous nucleic acids was not required. Instead, the upregulation of the IFN-I pathway could be explained solely by a cooperative effect of prolonged activation of NF-*κ*B and JNK pathways. Both pathways are known to converge on IFNβ enhancer and recruit coactivators and chromatin-remodeling proteins to form an enhanceosome[55, 56]. JNK is activated in response to oxidative, proteotoxic, metabolic and other challenges and is an important part of the cellular defense strategy against stress[57, 58].Whereas transient JNK activation is adaptive, prolonged JNK activation is known to contribute to pro-apoptotic transition, which, as we show, in TNF-stimulated *sst1*-susceptible macrophages occurs via type I IFN pathway upregulation and PKR activation.

We propose that stress-induced IFN-I pathway escalation mechanistically represents an intermediate step between a low grade constitutive IFN-I pathway activity that plays homeostatic role[59] and a full blown activation downstream of nucleic acid recognition receptors mediated by IRF3. Normally, it plays an adaptive role. For example, upregulation of JNK by proteotoxic stress induced by TNF may lead to transient activation of the IFNβ – PKR – ISR axis to limit global protein biosynthesis by inhibiting cap-dependent translation. However, un-resolving proteotoxic stress and persistent ISR observed in susceptible macrophages lead to further escalation of JNK activity, corresponding increase in IFNβ production and upregulation of interferon-stimulated genes PKR, Rsad2 and Ch25h, whose products inhibit protein biosynthesis, mitochondrial function and lipogenesis, respectively[54, 60, 61]. The 25-hydrocholesterol produced by the Ch25h enzymatic activity can further increase ISR[62] and amplify inflammatory cytokine production[63]. Moreover, by limiting cholesterol biosynthesis it can sustain elevated IFN-I signaling[64]. Previously, PKR has been shown to stimulate JNK activity in macrophages [42]. Possibly, this occurs via translational arrest, since other protein synthesis inhibitors also activate stress kinases JNK and p38 via a mechanism known as the “ribotoxic stress response”[65, 66]. Thus, it appears that at certain levels JNK, IFNβ and PKR may form a feed forward stress response circuit, locking TNF-stimulated sst1-susceptible macrophages in a state of unrelenting stress and, eventually, leading to suppression of essential metabolic pathways, IFN-I-dominated hyper-inflammatory response and macrophage damage.

Numerous studies reveal a role of the type I interferon (IFN-I) pathway in the immunopathology caused by intracellular bacteria, including Mtb[67-70], chronic viral infections[71-73] and autoimmunity (reviewed in[74-76]). Our studies reveal a mechanism of stress-mediated IFN-I pathway upregulation that makes macrophages less resilient to subsequent infection with intracellular bacteria and is associated with immunopathology in vivo. Obviously, this susceptibility-associated mechanism represents an attractive therapeutic target. On a cautionary note, however, elements of this pathway may represent imperfect, but necessary, backup strategy of stress adaptation, and their inhibition may be detrimental[11]. Thus, we observed that inhibition of HSF1 and JNK increases expression of stress markers, while blocking the IFN-I signaling increases replication of intracellular bacteria in TNF-primed susceptible macrophages. In contrast, inhibiting transcriptional activity of c-Myc, which is upstream of stress initiation in the sst1S phenotype, improved both macrophage survival and the bacterial control in vitro. We hypothesize that in vivo, Myc downregulation is a part of macrophage reprogramming within inflammatory milieu that allows balancing metabolism and pre-adaption to pathogen encounter.

Despite its focus on effects of a single genetic locus, our study reveals a generalizable paradigm in host - pathogen interactions. Traditionally, susceptibility to a specific pathogen is viewed as immune defect associated with inability to mount appropriate effector responses. Therefore, studies have been primarily focused on molecules involved in pathogen recognition and immune effector functions. A wealth of information about those molecules and associated essential mechanisms of host immunity has been obtained studying extreme susceptibility to infections in humans and knockout mice (reviewed in [77-79]. Indeed, avoidance or suppression of host recognition and effector mechanisms by a microbe are strategies essential for establishing infections. Our study demonstrates that subsequent disease progression in immune competent hosts may depend on locally induced macrophage susceptibility that emerge gradually within inflammatory tissue due to an imbalance of macrophage growth, differentiation and stress responses prior to contact with microbes. This explains how successful pathogens may exploit this regulatory failure to bypass mechanisms of resistance in otherwise immune competent hosts. This strategy would ensure survival of both the host and the pathogen and facilitate successful transmission of the later[80]. The origins of aberrant macrophage responses within inflammatory milieu may encompass genetic and non-genetic causes, such as co-infections, metabolic diseases and senescence. Since the *sst1*-susceptible phenotype in mice closely resembles pathology of human TB, we propose that environmental exposures, metabolic stressors, senescence and co-infections linked to TB risk or severity may also activate this nascent mechanism of infection susceptibility and immunopathology in humans. This concept suggests a novel disease-modifying therapeutic strategy focusing on correcting aberrant response to TNF rather than blocking this essential mediator of host resistance.

## MATERIALS & METHODS

### Reagents

Recombinant mouse TNF was from Peprotech and recombinant mouse IL-3 was from R&D. Mouse monoclonal antibody to mouse TNF (MAb; clone XT22) was from Thermo scientific and isotype control and mouse monoclonal antibody to mouse IFNb (Clone: MAR1-5A3) was from eBiosciences. BAY 11-7082, Phenylbutyrate sodium (PBA), rapamycin were from Enzo Life sciences. SB203580, SP600125 and C16 were obtained from were from calbiochem. JQ1, Flavopiridol, 10058-F4 were from Tocris. ISRIB, poly I:C, LPS from E. coli(055:B5), Triptolide and BHA were obtained from Sigma. BX-795 was from Invivogen. RHT was kindly provided by Aaron Beeler, BU CMD and Chemistry Department.

### Animals

C57BL/6J and C3HeB/FeJ inbred mice were obtained from the Jackson Laboratory (Bar Harbor, Maine, USA). The C3H.B6-sst1, C3H.scid and C3H.B6-sst1, scid mouse strains were generated in our laboratory as described previously[17, 20, 81]. The B6.C3H-sst1(B6J.C3-sst1C3HeB/FejKrmn) mice were created by transferring the *sst1* susceptible allele on the B6 (C57BL/6J) genetic background using twelve backcrosses. All experiments were performed with the full knowledge and approval of the Standing Committee on Animals at Boston University in accordance with relevant guidelines and regulations (IACUC protocol number AN15276).

### BMDMs culture and infection with F. tularensis LVS

Isolation of mouse bone marrow and culture of BMDMs were carried out as previously described[82]. TNF-activated macrophages were obtained by culture of cells for various times with recombinant mouse TNF (10 ng/ml). F. tularensis live vaccine strain (LVS) were grown in Brain-Heart Infusion broth overnight, harvested and then diluted in media without antibiotics to get the desired MOI. BMDM were seeded in tissue culture plates. Cells were treated with TNF and inhibitors were added after 4hrs of TNF addition. After 24hrs of treatment, cells were infected at indicated MOI. The plates were then centrifuged at 500× g for 15 minutes and incubated for 1hr at 37°C with 5% CO_2_. Cells were then washed with fresh media, and incubated for 45 min at 37°C with media containing gentamicin (50 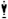 g/mL) to kill any extracellular bacteria. Cells were washed again and cultured in presence of inhibitors and TNF in DMEM/F12 containing 10% FBS medium without antibiotic at 37°C in 5% CO_2_ for 24hrs.

### Bacterial DNA isolation and quantification

For isolation of LVS DNA from 96-well plates, cells were lysed with 50 ul of 250mM NaOH, 0.2 mM EDTA and kept at RT for 10 mins. The samples were then heated for 45 mins at 95°C and neutralized with 50 ul 40mM Tris-HCL, pH 8.0. DNA was then isolated by purification with magnetic beads. TaqMan PCR conditions were carried out according to previous studies(**ref**). All PCR reactions were performed in a final volume of 20ul and contained Taqman Environmental mastermix (Applied Biosystems) at a 1X final concentration, probe (250nM), and primers (450nM). The primers used were FopAF: ATCTAGCAGGTCAAGCAACAGGT, FopAR: GTCAACACTTGCTTGAAC-ATTTCTAGATA, and the probe FOPAP: CAAACTTAAGACCACCACCCACATCCCAA. Thermal cycling conditions were 50°C for 2 min, 95°C for 10 min, 45 cycles at 95°C for 10 s and 60°C for 30 s, and then 45°C for 5 min.

### Immunoblotting

To monitor the Ipr1 protein levels we have developed Ipr1 peptide-specific rabbit polyclonal antibodies, which recognized the Ipr1 protein of predicted length on Western blots (ref Sc Report paper). BMDM’s were subjected to treatments specified in the text. Nuclear extracts were prepared using the nuclear extraction kit from signosis. Whole cell extracts were prepared by lysing the cells in RIPA buffer supplemented with protease inhibitor cocktail and phosphatase inhibitor I and III (Sigma). Equal amounts (30 μg) of protein from whole-cell extracts was separated by SDS-PAGE and transferred to PVDF membrane (Millipore). After blocking with 5% skim milk in TBS-T buffer [20 mMTris–HCl (pH 7.5), 150 mM NaCl, and 0.1% Tween20] for 2 hour, the membranes were incubated with the primary antibody overnight at 4 °C. Bands were detected with enhanced chemiluminescence (ECL) kit (Perkin Elmer). Stripping was performed using WB stripping solution (Thermo scientific). The loading control β-actin (Sigma, 1:2000) was evaluated on the same membrane. The Ipr1-specific rabbit anti-serum was generated by Covance Research Products, Inc. (Denver, CO, USA)(1:500) as described previously[82]. The Ipr1 monoclonal antibodies were generated using Ipr1 peptides from Abmart. ATF4, ATF3, Gadd34, c-Myc, Daxx, p21, PKR and phospho-PKR antibodies were obtained from Santacruz biotechnology. IRF1, IRF3 (1:1000), p38, p-p38, JNK, p-JNK antibodies were obtained from Cell signaling. Hspa1a (1:1000) antibody was obtained from R&D. ß-actin (1:2000) was obtained from Sigma. Bim and DHCR7 were obtained from Abcam.

### RNA Isolation and quantitative PCR

Total RNA was isolated using the RNeasy Plus mini kit (Qiagen). cDNA synthesis was performed using the SuperScript II (Invitrogen). Real-time PCR was performed with the GoTaq qPCR Mastermix (Promega) using the CFX-90 real-time PCR System (Bio-Rad).Oligonucleotide primers were designed using Primer 3 software (**Supplementary Table S1)** and specificity was confirmed by melting curve analysis. Thermal cycling parameters involved 40 cycles under the following conditions: 95 °C for 2 mins, 95 °C for 15 s and 60 °C for 30 s. Each sample was set up in triplicate and normalized to RPS17 or 18S expression by the DDCt method.

### Immunofluorescence microscopy

Cells were fixed with 4% paraformaldehyde for 15 min at RT, cells were permeabilised with 0.25% Triton-X for 30 min and then blocked for 20 min with goat-serum (2.5%). Cells were incubated with primary antibodies [mouse monoclonal antibodies against J2 (1:3000), overnight at 4 °C in 2.5% goat serum, and incubated with Alexa Fluor 488-conjugated donkey anti-mouse IgG (excitation/emission maxima ∼ 490/525 nm) (1:1000, Invitrogen) secondary antibody for 2 hrs. Images were acquired using Leica SP5 confocal microscope. All images were processed using Image J software.

### Hoechst/PI Staining Method for cell cytotoxicity

For cell viability assays BMDM were plated in 96 well tissue culture plates (12000 cells/well) in phenol-red free DMEM/F12 media and subjected to necessary treatments. Hoechst (Invitrogen, 10 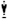 M) and PI (Calbiochem, 2 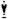 M) were added. The plates were kept at 37 °C for 15 min and read in the celigo cell cytometer. The % of total and dead cells was calculated for each treatment.

### Transcription factor profiling analysis

Each array assay was performed following the procedure described in the TF activation profiling plate array kit user manual (Signosis, Inc, FA-001). 10 ug of nuclear extract was first incubated with the biotin labeled probe mix at room temperature for 30 min. The activated TFs were bound to the corresponding DNA binding probes. After the protein/DNA complexes were isolated from unbound probes, the bound probes were eluted and hybridized with the plate pre-coated with the capture oligos. The captured biotin-labeled probes were then detected with Streptavidin– HRP and subsequently measured with the TECAN microplate reader.

### Gel shift assay

The nuclear extracts with 12 h of TNF treatment in sst1R and sst1S were chosen for gel shift assay analysis with EMSA kits (Signosis Inc). The TF DNA binding probe sequences are listed below.

1. AP1: CGCTTGATGACTCAGCCGGAA

2. c-Myc: AGTTGACCACGTGGTCTGGG

The sequences that we used as probes for gel shift assay are identical to those we used as the probe mix for TF activation profiling array assay. 5ug nuclear extracts were incubated with 1× binding buffer and biotin-labeled probe for 30 min at room temperature to form protein/DNA complexes. The samples were then electrophoresed on a 6 % polyacrylamide gel in 0.5 % TBE at 120 V for 45 min and then transferred onto a nylon membrane in 0.5 % TBE at 300 mA for 1 h. After transfer and UV cross-linking, the membrane was detected with Streptavidin–HRP. The image was acquired using a FluorChem imager (Alpha Innotech Corp).

### siRNA knockdown

Gene knockdown was done using GenMute (SignaGen) and Flexitube Genesolution siRNAs from Qiagen. All star negative control siRNA (SI03650318) from Qiagen was used as a negative control. sst1S and sst1R BMDMs were seeded into 6-well plates at a density of 2.5 × 10^5^ per well and grown as mentioned before. Shortly before transfection, the culture medium was removed and replaced with 1 ml complete medium, and the cells were returned to normal growth conditions. To create transfection complexes, 15 nM siRNA (pool of 4 siRNAs) in 1× GenMuteBuffer (total 500 ml) was incubated with 1.5 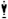 l of Genemute transfection reagent for 15-20 minutes at room temperature. The complexes were added drop-wise onto the cells. Cells were incubated with the transfection complexes for 24 hours at 37° in 5% CO2. After 24hrs cells were washed to remove siRNA and replaced with fresh media. TNF (10ng/ml) was added for 24hrs and BMDMs were harvested as outlined below. siRNA pools included: Irf1 (GS16362), Irf3 (GS54131), Irf7(GS54123)

### ELISA

Supernatants were collected from mouse macrophages after 24hrs of stimulation with TNFa or poly IC. IFNb was measured using the mouse IFN-b ELISA kit from pbl Assay Science. ELISAs were done as recommended by the manufacturer.

## Acknowlegement

This work was sponsored by R01 HL133190 and R01 HL126066 to IK. The authors are grateful to Drs. Robert Silverman and Benjamin Wolozin for helpful discussions, and to Adam Gover and Somak Ray for the analyses of microarray and proteomics data, respectively.

The authors declare no competing interests.

