## Supplementary Materials for "Increased susceptibility to intracellular bacteria and necrotic inflammation driven by a dysregulated macrophage response to TNF"

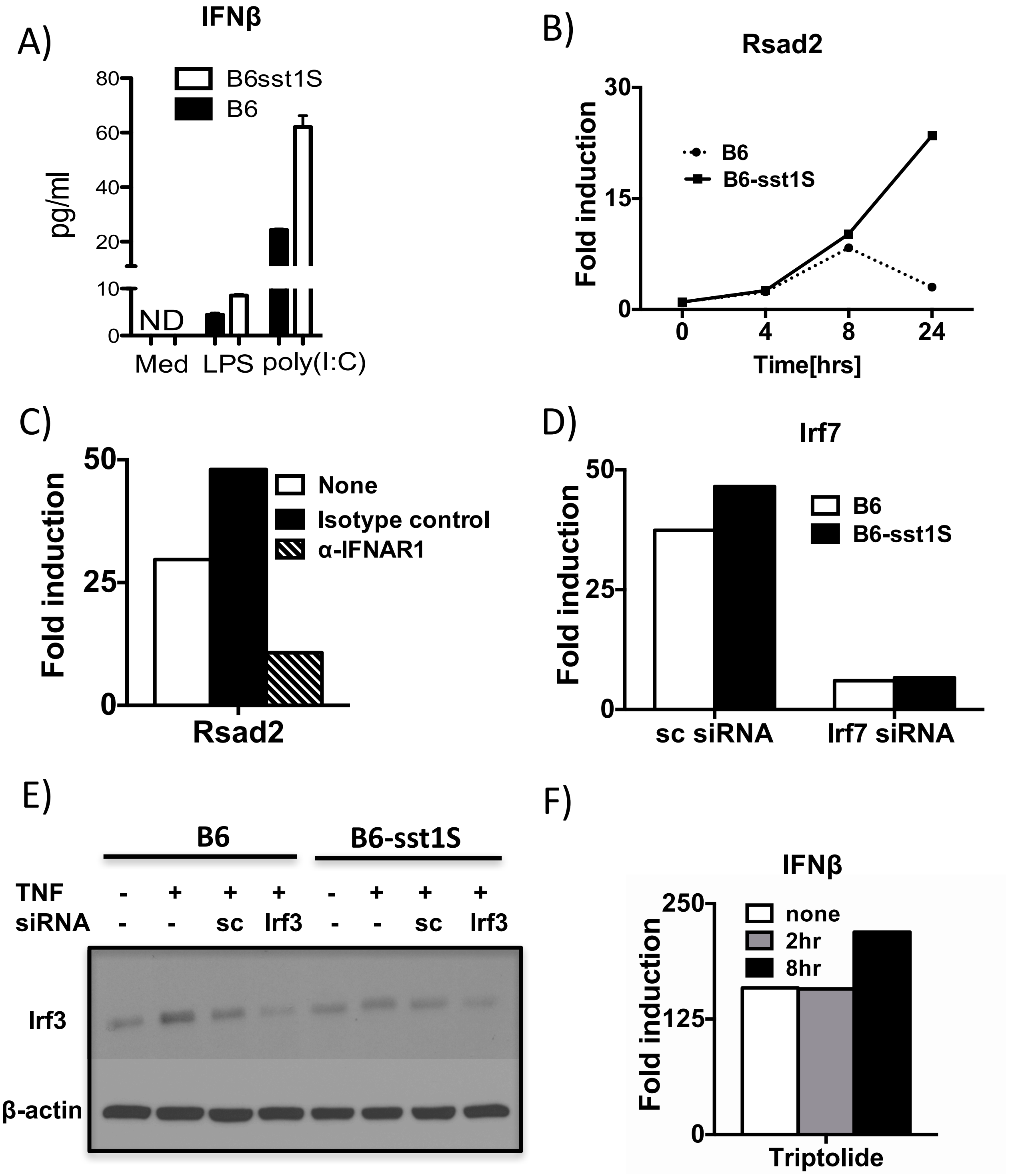
A)
IFNβ
B6sst1S
B6
pg/ml
80
60
40
20
10
0
ND
Med
LPS
poly(I:C)
B)
Rsad2
B6
B6-sst1S
Fold induction
30
15
0
0
4
8
24
Time[hrs]
C)
None
Isotype control
α-IFNAR1
Fold induction
50
25
0
Rsad2
D)
Irf7
B6
B6-sst1S
Fold induction
50
25
0
sc siRNA
Irf7 siRNA
E)
B6
B6-sst1S
TNF - + + + - + + +
siRNA - - sc Irf3 - - sc Irf3
Irf3
β-actin
F)
IFNβ
none
2hr
8hr
Fold induction
250
125
0
Triptolide

**Suppl. Fig 1**

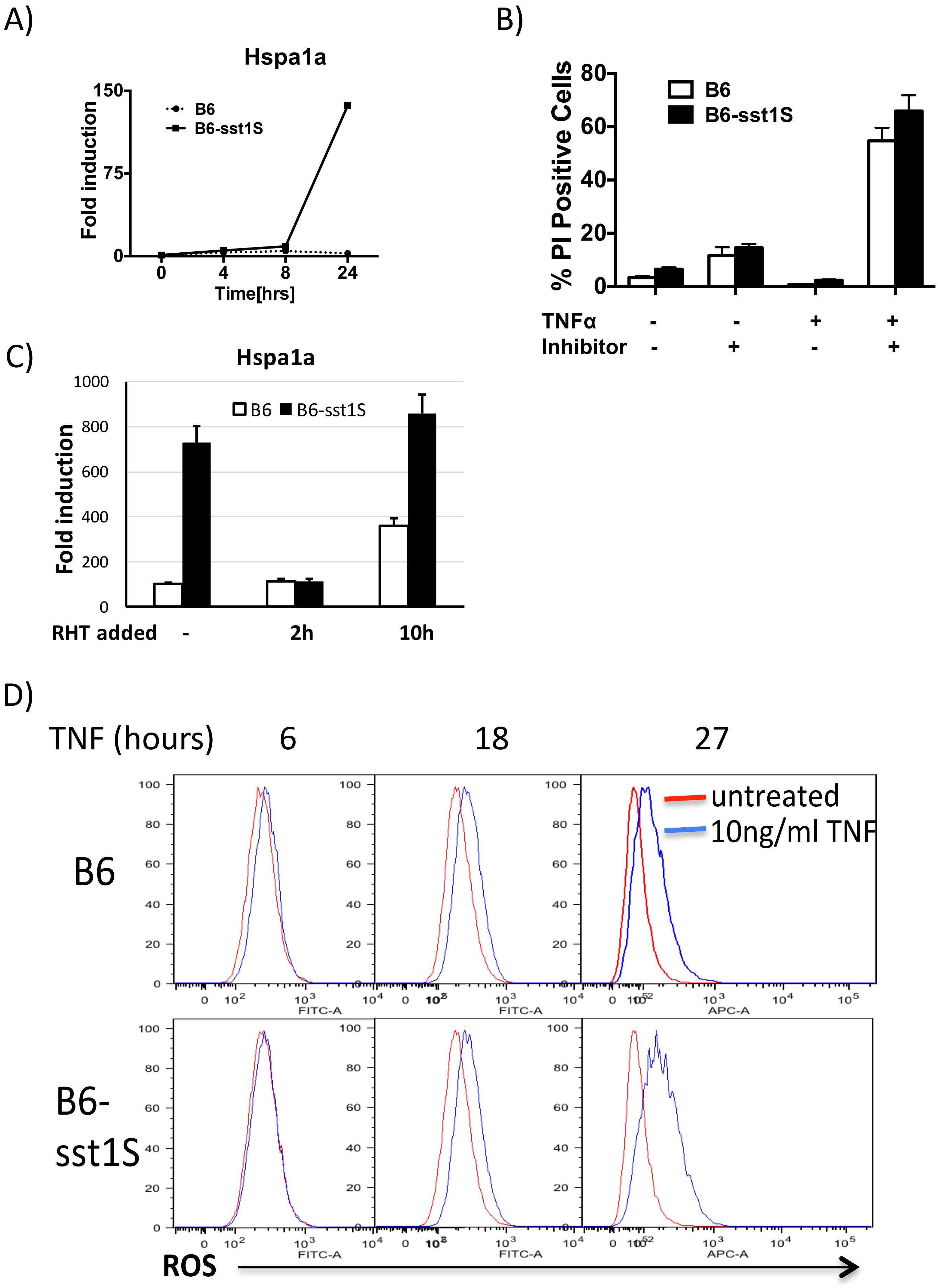

A)
Hspa1a
B6
B6-sst1S
Fold induction
Time[hrs]
B)
% PI Positive Cells
B6
B6-sst1S
TNFα - - + +
Inhibitor - + - +
C)
Hspa1a
B6 B6-sst1S
Fold induction
RHT added - 2h 10h
D)
TNF (hours) 6 18 27
B6
B6-sst1S
untreated
10ng/ml TNF
FITC-A
APC-A
ROS


**Suppl. Fig 2**

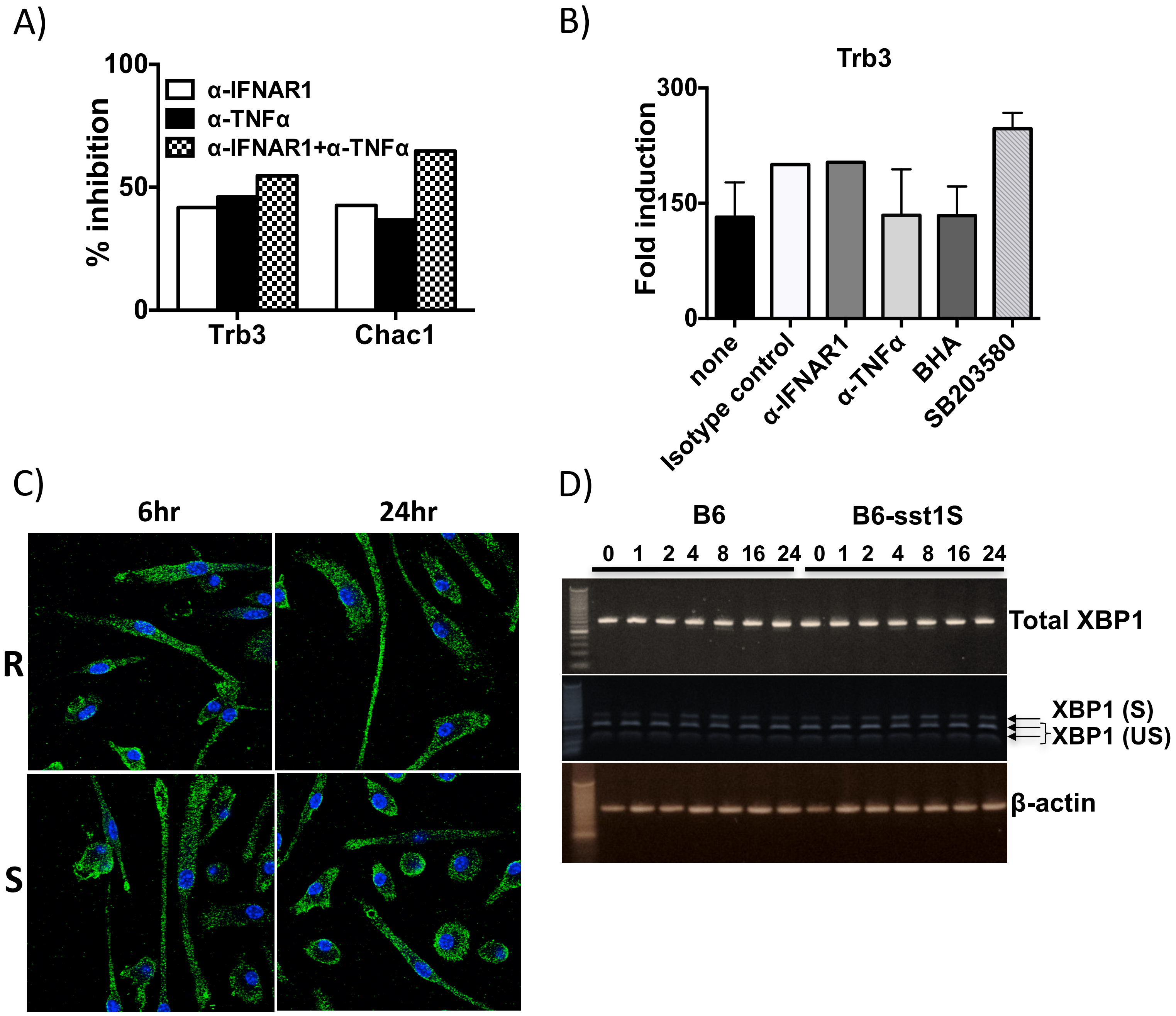
A)
100
50
0
% inhibition
α-IFNAR1
α-TNFα
α-IFNAR1+α-TNFα
Trb3
Chac1
B)
Trb3
300
150
0
Fold induction
none
Isotype control
α-IFNAR1
α-TNFα
BHA
SB203580
C)
6hr
24hr
R
S
D)
B6
B6-sst1S
0 1 2 4 8 16 24 0 1 2 4 8 16 24
Total XBP1
XBP1 (S)
XBP1 (US)
β-actin

Suppl. Fig 3

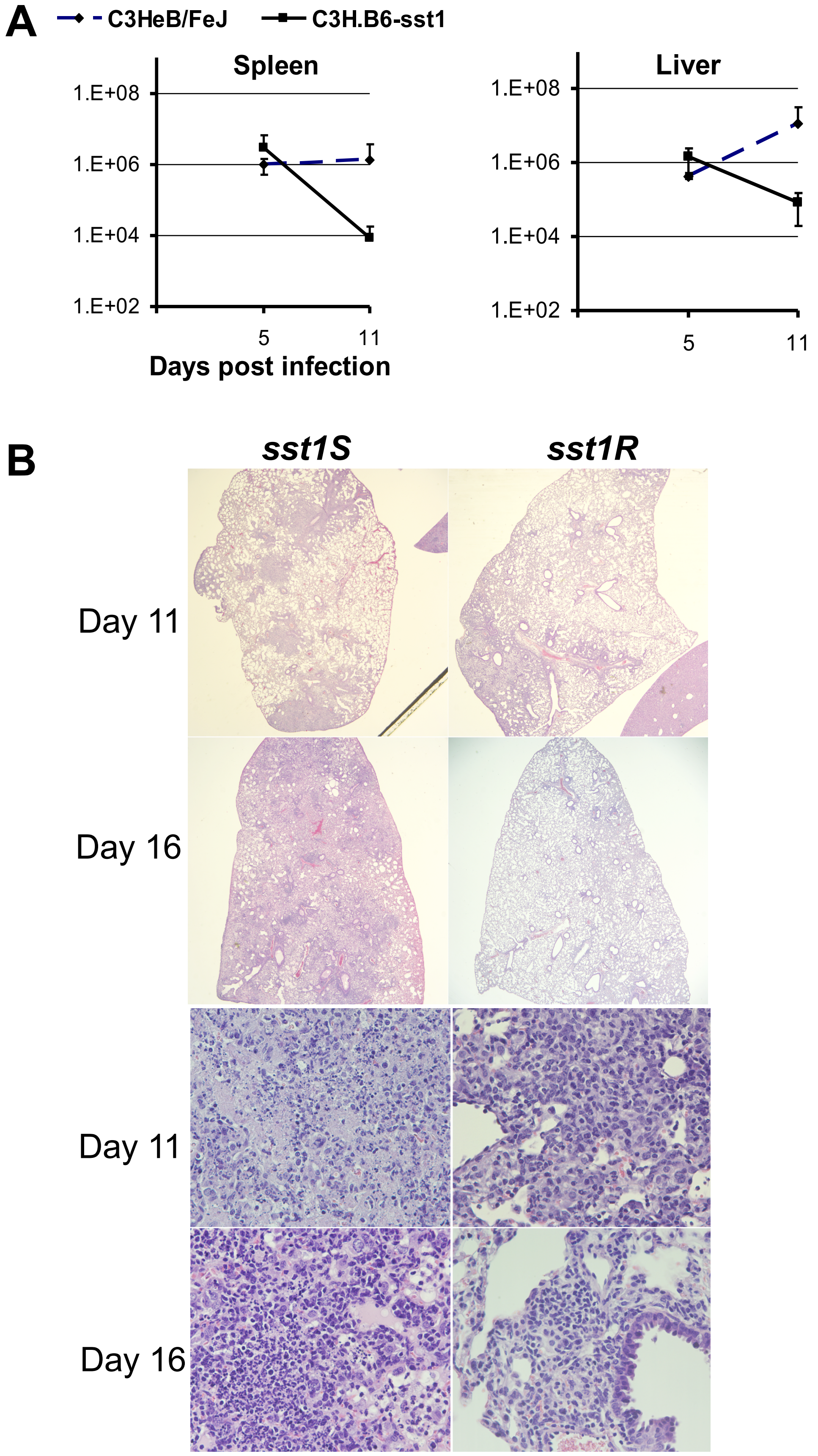
A
C3HeB/FeJ
C3H.B6-sst1
Spleen
Liver
1.E+08
1.E+06
1.E+04
1.E+02
5
11
Days post infection
B
sst1S
sst1R
Day 11
Day 16
Day 11
Day 16

**Suppl. Fig 4**

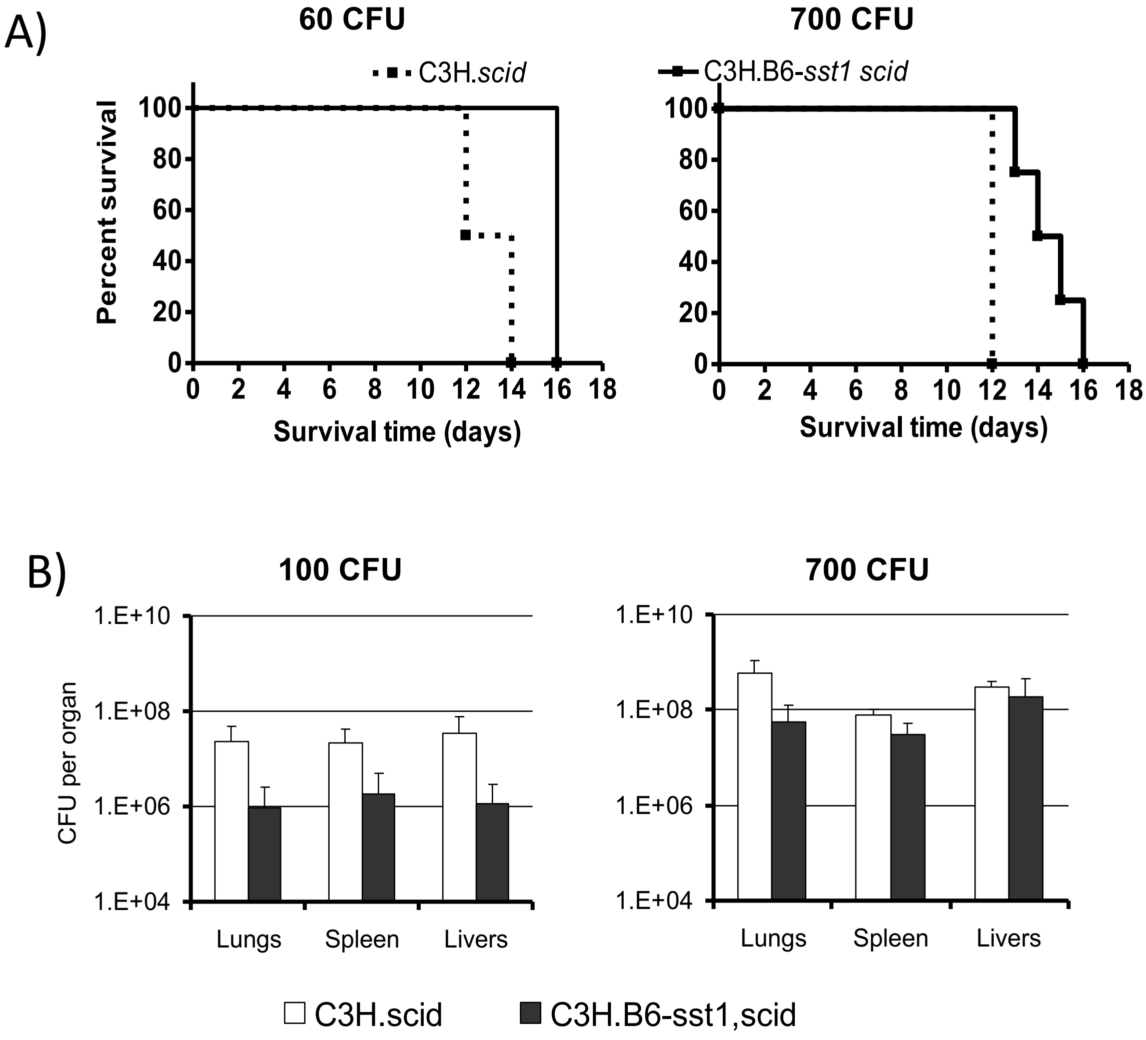
A)
60 CFU
700 CFU
C3H.*scid*
C3H.B6-*sst1 scid*
Percent survival
100
80
60
40
20
0
0 2 4 6 8 10 12 14 16 18
Survival time (days)
B)
100 CFU
700 CFU
CFU per organ
1.E+10
1.E+08
1.E+06
1.E+04
Lungs
Spleen
Livers
C3H.scid
C3H.B6-sst1,scid

**Suppl. Fig 5**

**Supplementary Figure 1. A)** IFNβ concentrations in supernatants of B6wt and B6-sst1S BMDM treated with 100ng/ml LPS and 1ug/ml poly IC for 24h, as determined by ELISA. **B**) Time course of Rsad2 mRNA expression in B6wt and B6-sst1S BMDM treated with 10ng/ml TNF (representative of two independent experiments). **C**) Inhibition of Rsad2 mRNA expression in TNF-stimulated B6-sst1S BMDM using α-IFNAR1 blocking antibodies. TNFα (10ng/ml) and blocking antibodies or isotype control were added simultaneously for the duration of experiment (24h). **D**) Validation of Irf7 knockdown in B6wt and B6-sst1S BMDM. Irf7 mRNA expression was quantified using qRT-PCR, normalized to expression of 18S rRNA and presented relative to expression in untreated cells (set as 1). The data are representative of two independent experiments. **E**) Validation of IRF3 knockdown using immunoblotting with IRF3-specific antibodies (sc – scrambled control; Irf3 – specific siRNA). **F**) IFNβ mRNA expression in B6-sst1S BMDM treated with 10ng/ml TNF for 18hrs in the presence of 500 nM Triptolide added at 2 or 8 h of TNF treatment (no significant difference between treatment groups and control).

**Supplementary Figure 2. A)** Timecourse of Hspa1a mRNA expression in B6 and B6-sst1S BMDM stimulated with TNF (10ng/ml). **B)** Synergistic effect of TNF and HSF1 inhibitor KRIB11 on macrophage death. B6 and B6-sst1S BMDM were treated with 10ng/ml TNFα for 24 hrs in presence or absence of 10uM KRIB11. Cell death was assessed using automated microscopy as % of PI positive cells. Data represents results from two independent experiments. **C)** Effect of the HSF1 and translation inhibitor rohitinib (RHT) on Hspa1a mRNA expression in B6-sst1S BMDM treated with 10ng/mL of TNF for 24 hrs. RHT (300uM) was added at 2 and 10 h of TNF stimulation. Gene expression data are normalized to expression of 18S mRNA and presented relative to expression in untreated cells (set as 1). Results represent data from three independent experiments. **D)** ROS production in B6wt and B6-sst1S BMDM stimulated with 10ng/ml TNF$\alpha$ for the indicated times. FACS analysis was performed using DCFDA (dichlorofluorescindiacete) probe (Abcam).

**Supplementary Figure. 3 A)** Trb3 and Chac1mRNA expression in B6-sst1S BMDM treated with 10ng/ml TNF$\alpha$ for 18 h in the presence of either $\alpha$-IFNAR1 (10 ug/ml), $\alpha$-TNF$\alpha$ (10 ug/ml) or their combination. **B**) No effect of late IFNAR1 and TNF blockade or ROS (BHA) and p38 (SB203580) inhibitors on Trb3 mRNA expression in B6-sst1S BMDM stimulated with TNF for 16 h. All inhibitors were added at 12 h of TNF treatment. **C**) Staining with dsRNA-specific J2 antibodies of B6wt and B6-sst1S BMDM treated with 10ng/ml TNF for 6 and 24 h. Cells were stained with J2 antibody (green); nuclei are counterstained with DAPI (blue). All microscopic images represent data from two independent experiments. **D**) Timecourse analysis of XBP-1 mRNA expression and splicing in TNF-stimulated B6wt and B6-sst1S BMDM. Total RNA from TNF stimulated B6wt and B6-sst1S BMDM were amplified by RT-PCR as described in Methods. After digestion with Pst1, the PCR products were subjected to 2% agarose gel electrophoresis. The PCR products of spliced XBP1(S) remained intact, whereas the PCR products of unsplicedXBP1(U) mRNA were cut into two fragments as indicated by arrows.

**Supplementary Figure 4. A**) The kinetics of *F.t.* LVS growth in spleens and livers of the *sst1*$^{R}$ and *sst1*$^{S}$ mice after the aerosol infection with 1600 CFU of *F.t.* LVS. Four mice per group were used permouse strain at each time point. **B**) Histopathology of the lungs of *sst1*$^{S}$ and *sst1*$^{R}$mice 11 and 16 days after aerosol infection with 1600 CFU of *F.t.* LVS. H&E staining, magnification 20X (upper panels) and 200X (lower panels). esrAS

**Supplementary Figure 5. A)** Survival of the sst1 congenic *scid* mice after aerosol infection with 60 or 700 CFU of *F.t.* LVS by aerosol.C3H.B6-*sst1,scid*(*sst1*$^R$ - solid line) and C3H.*scid*(*sst1*$^S$ - dashed line)mouse strainswere used in this experiment.Four to six mice per group were used for each strain. **B**) *F.t.* LVS burdens in the organs of the *sst1*$^R$ and *sst1*$^S$ mice (C3H.B6-*sst1,scid* and C3H.*scid*, respectively) 11 days after aerosol infection with 100 or 700 CFU of F. LVS. Four mice per group were used at each time point.

**Microarray analysis**

BMDM from sst1R and sst1S mice were treated with 10 ng/ml TNF$\alpha$ for 18hrs. Total RNA was isolated using RNeasy Plus Mini kit (Qiagen) according to manufacturer's instructions. Biotin labeling was performed using the Ambion WT Expression Kit (Life Technologies, Grand Island, NY) according to the manufacturer's protocol, followed by the GeneChip WT Terminal Labeling and Controls Kit (Affymetrix, Santa Clara, CA). The labeled, fragmented DNA was hybridized to the Affymetrix GeneChip Mouse Gene 2.0 ST Array for 18 hours in a GeneChip Hybridization oven 640 at 45$^{o}$C with rotation (60 rpm). The hybridized samples were washed and stained using an Affymetrix fluidics station 450 with streptavidin-R-phycoerythrin (SAPE) and the signal was amplified using a biotinylated goat anti-streptavidin antibody followed by another SAPE staining (Affymetrix Hybridization, Washing and Staining Kit). After staining, microarrays were immediately scanned using an Affymetrix GeneArray Scanner 3000 7G Plus. CEL files were normalized to produce gene-level expression values using the implementation of the Robust Multiarray Average (RMA) in the affy package (version 1.36.1) included within in the Bioconductor software suite (version 2.11) and an Entrez Gene-specific probeset mapping (version 17.0.0) from the Molecular and Behavioral Neuroscience Institute (Brainarray) at the University of Michigan. Array quality was assessed by computing Relative Log Expression (RLE) and Normalized Unscaled Standard Error (NUSE) using the affyPLM Bioconductor package (version 1.34.0). Analyses of variance were performed using the f.pvalue function in the sva package (version 3.4.0). Pairwise differential gene expression was assessed by performing Student's t test on the coefficients of simple linear models computed using the lmFit function in the limma package (version 3.14.4). Correction for multiple hypothesis testing was accomplished using the Benjamini-Hochberg false discovery rate (FDR). Human homologs of mouse genes were identified using HomoloGene (version 67). All microarray analyses were performed using the R environment for statistical computing (version 2.15.1).

**Gene Set Enrichment Analysis (GSEA)**

GSEA (version 2.0.13) was used to identify biological terms, pathways and processes that were coordinately up- or down-regulated within each pairwise comparison. The Entrez Gene identifiers of the human homologs of the genes interrogated by the array were ranked according to the t statistic computed for the TNF$\alpha$ versus untreated comparison within each strain. Mouse genes without a human homolog were removed, and t statistics for multiple mouse genes with the same human homolog were averaged prior to ranking. The resulting ranked lists were each used to perform a pre-ranked GSEA analysis (default parameters with random seed 1234) using the Entrez Gene versions of the Biocarta, KEGG, Reactome, Gene Ontology (GO), and transcription factor and microRNA motif gene sets obtained from the Molecular Signatures Database (MSigDB), version 4.0.

**Sample preparation for proteomics**

sst1R and sst1S BMDM were treated with 10ng/ml TNF$\alpha$ for 24 hrs. Media was removed and cells were rinsed with dPBS. 150-200 uL of the lysis buffer (8 M urea, 2 M guanidinium chloride, 5 mM DTT in 100 mM TEAB (pH 8.0) freshly prepared was added to cover the whole area of the bottom of the well. Phosphatase and protease inhibitor cocktail was added into the lysis buffer at a final conc of 1%. Cells were scraped and the plate was incubated for 20 minutes while rocking the plate on a rocking shaker at a low speed at room temperature (to avoid crystallization of urea) to enable efficient lysis. The lysate was transferred into a clean low retention Eppendorf tube. Cell debris was removed by centrifuging the resulting lysate at 14,000g for 10 min at 4$^{o}$ C. The supernatant was transferred to a fresh, low retention Eppendorf tube pre-washed with MeOH and dried and stored at -80$^{o}$C.

### Analysis of XBP1 mRNA Splicing by IRE1

1 ug RNAs were reverse transcribed using Oligo(dT) primer (Invitrogen), and amplified using sense primer mXBP1.3S (5′-AAACAGAGTAGCAGCGCAGACTGC-3′) and antisense primer mXBP1.2AS (5′-GGATCTCTAAAACTAGAGGCTTGGTG-3′) to generate cDNA product encompassing the IRE1 cleavage sites as previously described (12). The unspliced and spliced mRNAs generate 480- and 454-bp cDNA products, respectively. These fragments were further digested by PstI to check whether a PstI restriction site was lost after IRE1-mediated splicing of mRNA. The cDNA fragments were resolved on 2% agarose gel. cDNA products from the unspliced mRNA yielded two short fragments (289 and 191 bp) after digestion.

### Infection of mice with Francisella tularensis *Live Vaccine Strain* (F. LVS)

F. LVS (ATCC 29684) was grown in modified Mueller-Hinton broth with Isovitalex (BD Biosciences), harvested and frozen at−80°C in 1ml aliquots ($10^{10}$ cfu/ml) in the presence of 10% (v/v) sucrose. Bacteria were defrosted, inoculated into fresh Mueller-Hinton broth and grown for a maximum of 8 hours at 37°C in an orbital incubator. Mice were exposed to aerosolized F. LVS in a 'nose-only' CH Technologies in-hood aerosol system with a SLAG aerosol generator [25] for 20-40 minutes at various dilutions of bacteria depending on desired level of infection. Infections were performed inside a Bio-Safety Level 3 facility. Mice were euthanized by halothane anaesthesia at various timepoints post-infection. Organs were homogenized in PBS and serial tenfold dilutions were cultured on Mueller-Hinton (Difco) plates enriched with horse serum (Hybrid), Isovitalex (BD Bioscience), Glucose (Sigma) and Ferrous oxide (Sigma) for 72 hours at 37°C.

### Histopathology

Mice were euthanized with halothane and necropsied. The lungs, livers and spleens of infected mice were removed, fixed immediately by immersion in 10% neutral buffered formalin for 24 hours, and processed by paraffin embedding. These tissues were sectioned and stained with hematoxylin and eosin (H&E) by standard procedure at the Harvard Rodent Histopathology Core Facility.

**Isolation of lung cells**

Isolation of cells from lungs was performed as described (1). Briefly, mice were euthanized by s.c. injection of sodium pentobarbital (100μl with 64.8 mg/ml), and suspensions of lung cells were prepared individually. Blood vessels were washed out with 20 ml of PBS containing heparin (10 U/ml), and repeated bronchoalveolar lavage was performed via canulated trachea using 5 ml of PBS. Lung tissue was sliced into 1- to 2-$mm^3$ pieces and incubated at 37°C in L-15 medium containing 5% FCS, kanamycin (0.05 μg/ml), 10 mM HEPES (all from HyClone), 150 U/ml collagenase IV, and 50 U/ml DNase I (Sigma-Aldrich). A total volume of 5 ml of digestion medium was used to digest lungs from each mouse during incubation in a shaker for 90 min. Single-cell suspensions were obtained by vigorous pipetting and filtration trough 100 μm cell strainer. Cells were washed 3 times with PBS supplemented with 2% FCS and counted. The viability of cells, as determined by trypan blue exclusion, was >95%.

**Flow cytometry**

3-5 $\times 10^5$ lung cells were washed twice in PBS containing 0.01 % $NaN_3$ and 0.5 % BSA, and incubated for 15 min at $4^0$C in the presence of CD16/CD32 mAbs (clone 2.4G2, BD Biosciences, San Diego, CA) to block Fc-receptors. Cells were then double-, triple-, or quadruple-stained with directly conjugated antibodies, according to the manufacturer's instructions. Specific monoclonal antibodies for mouse CD4 (L3T4 clone RM4-5, rat IgG2a), CD8 (Ly2 clone 53-6.7, rat IgG2a), CD49b (clone DX5, rat IgM), Ly6C (clone AL-21, rat IgM), Ly6G (clone 1A8, rat IgG2a) were purchased from BD Biosciences. Monoclonal antibody specific for mouse F4/80 (clone BM8, rat IgG2a) was purchased from Caltag Laboratories (Burlingame, CA) as direct conjugate to PE. Stained cells were washed twice, fixed with 2% paraformaldehyde and analyzed by flow cytometry, using FACSCalibur cytometer (BD) and BD-CellQuestPro (Beckton Dickinson) and FlowJo 4.5.9 (Tree Star, Inc., San Carlos, CA) software.

Monoclonal antibodies specific for mouse CD4 (L3T4 clone H129.19, rat IgG2a), CD8 (Ly-2 clone 53-6.7, rat IgG2a), CD49b (clone DX5, (and CD25 (IL-2R 🌡 chain clone 7D4, rat IgM) were purchased from BD (San Diego, CA) as direct conjugates to fluorescein isothiocyanate (FITC), phycoerythrin (PE), peridinin chlorophyll (PerCP) or biotin. Monoclonal antibody specific for mouse F4/80 (clone BM8, rat IgG2a) was purchased from Caltag Laboratories (Burlingame, CA) as direct conjugate to PE. Unseparated lung and lymph node cells were washed in PBS containing 1% BSA and 0.01%NaN3 and incubated for 10 minutes at 4$^{o}$C in the same buffer containing FC receptor blocking antibody (CD16/CD32, BD). After an additional wash, cells were triple stained for 30 minutes at 4$^{o}$C with directly or indirectly conjugated antibodies according to the manufacturer's instructions. Stained cells were washed three times in PBS containing 1% BSA and 0.01%NaN3, fixed in PBS containing 2% paraformaldehyde and analyzed by flow cytometry using FACSCalibur cytometer (BD) and CellQuest software (BD).

**Intracellular cytokine staining**

Stained for surface markers and fixed cells were permeabilized by washing 2 times in BD Perm/Wash buffer (BD) and incubated for 15 minutes in BD Perm/Wash buffer. Pellet cells were resuspended in BD Perm/Wash buffer containing PE-conjugated IFNγ antibody (BD, clone MXG1.2, rat IgG1) and incubated on ice for 30 minutes. After incubation cells were washed twice BD Perm/Wash buffer and resuspended in staining buffer prior to flow cytometric analysis (see above).

**Statistical analysis**

GraphPad Prizm 3.0 (GraphPad) software was used for the analysis. Comparison of bacterial loads was performed with Student's t-test. Results are presented as means ± s.d. The threshold for statistical significance was P, 0.05. Kaplan–Meier survival curves were generated and compared by using the log-rank test (GraphPad Prizm). The statistical significance was tested with P, 0.05 as the critical value with the Student–Newman–Keuls post-test to compare means between genetic backgrounds. Data are presented as means ± 95% confidence intervals for the mean.

### Supplementary Table S1: Primers used for real-time PCR amplification

| | | |
|---|---|---|
| *Ddit4* | CCTGCGCGTTTGCTCATGCC | GGCCGCACGGCTCACTGTAT |
| *ATF3* | CTGCAGAAAGAGTCGGAG | TGAGCCCGGACAATACAC |
| *Trib3* | GCAAAGCGGCTGATGTCTG | AGAGTCGTGGAATGGGTATCTG |
| *Chac1* | CCTGCTACCCTGCTC TTACCT | GAGCTTGGCTCCTCAGGTC |
| *Hspa1a* | GATTTGTTTTGCAGGACAGC | GGGGAGAGTCCAAACACAAA |
| *Hspa1b* | AAGAACGCGCTCGAGTCCTAT | TTGTCAGCCTCGCTGAGCTT |
| *Irf7* | CTGGAGCCATGGGTATGCA | AAGCACAAGCCGAGACTGCT |
| *Ipr1* | ACACTCCCTGTGACCTGTGG | GCCAATCTCCTGCCTCATTC |
| *Irf1* | CAGAGGAAAGAGAGAAAGTCC | CACACGGTGACAGTGCTGG |
| *18S* | TCAAGAACGAAAGTCGGAGGT | CGGGTCATGGGAATAACG |
| *ß-actin* | GTGGGCCGCTCTAGGCACCA | CGGTTGGCCTTAGGGTTCAGGG |
| *Rsad2* | AAGCTGAGGAGGTGGTGCAG | GAAAACCTTCCAGCGCACAG |
| *Ddit3* | GTCCCTAGCTTGGCTGACAGA | TGGAGAGCGAGGGCTTTG |
| *IFN*🚲 | ATGAGTGGTGGTTGCAGGC | TGACCTTTCAAATGCAGTAGATTC |
| *IP10* | GACGGGTCCGCTGCAACT | GCTTCCCTATGGCCCTCATT |
| *IL10* | CCAGTACAGCCGGGAAGACAATA | TGGCAACCCAAGTAACCCTTAAA |
| *RPS17* | TGTCGGGATCCACCTCAATG | CGCCATTATCCCCAGCAAG |
| *Ch25h* | CCATCTTTACCTTTCACGTGATTAC | CAGCCAAAGGGCACAAGTCT |
| *Fbxw7* | ACACGTTACGGGACACACTAATAGT | ACCACATGGATGCCATCAAAC |
| *Tnfrs12* | ATGGACTGCGCTTCTTGTC | CCAGAATGGGCCACAGTAG |
| *Bhlhe40* | CCAGGCCTCAACACCTCAGCTG | CCGAAGAGTCGAGGGACGAATG |
| *Bhlhe41* | AACATGGACGAAGGAATCCCTC | TAAGGCTGTTAGCGCTTTCAAG |
| *Trb1* | CCTCGAATATGGCAGCATTT | CGAGTCTCCTCACCCTTGTC |
| *MMP13* | AAGATGTGGAGTGCCTGATG | AAGGCCTTCTCCACTTCAGA |
| *XBP1* | GAACCAGGAGTTAAGAACACG | AGGCAACAGTGTCAGAGTCC |
